# Cell type-resolved connectomic reconstruction of the spinal dorsal horn reveals sensory neuron synaptic organization and modality-specific inhibition

**DOI:** 10.64898/2026.09.27.754691

**Authors:** Wangchu Xiang, Kevin Delgado, Qiyu Zhang, Josef Turecek, Adrian Holtrup, Ramin Khajeh, Tri Nguyen, J. Alex Bae, Akhilesh Halageri, Nico Kemnitz, Thomas Macrina, Dong Zhang, Sidrick Cameron, Borja Moya Llamas, Mohd Freezely Mazri, Julianne Ni, Ruairi Roberts, Alice Schildkamp, Laia Serratosa Capdevila, Aliya Norton, Jingyi Liu, Mingguan Liu, Sven Dorkenwald, David L. Paul, David D. Ginty, Wei-Chung Allen Lee

**Affiliations:** Department of Neurobiology, Harvard Medical School, 220 Longwood Avenue, Boston, MA, USA; Howard Hughes Medical Institute, Harvard Medical School, 220 Longwood Avenue, Boston, MA, USA; Zetta AI, Sherrill, NJ, USA; Aelysia LTD, Bristol, UK; Allen Institute for Brain Science, Seattle, WA, USA; FM Kirby Neurobiology Center, Boston Children’s Hospital, Boston, MA

## Abstract

Touch, temperature, chemical, and noxious stimuli are detected by more than a dozen peripheral sensory neuron subtypes that together mediate somatosensation. Somatosensory neurons form synapses in the spinal cord dorsal horn, the first site of central somatosensory processing, to engage the downstream circuitry that drives both spinal reflexes and ascending pathways for perception. Yet how sensory neuron synapses are organized into the spinal cord circuitry remains poorly understood. Here we present two connectomic tools for studying somatosensory circuits of the spinal cord dorsal horn: dSC1, a large-scale, synapse-resolved electron microscopy (EM) dataset of the mouse dorsal horn, and SPINE (SPinal cord Integrated NEuron classifier), a deep-learning pipeline that identifies major sensory neuron subtypes directly from EM images. Applying SPINE to the dSC1 dataset, we found marked heterogeneity in synaptic architecture across physiologically distinct sensory neuron subtypes: low-threshold mechanoreceptors form substantially more synapses and receive far more GABAergic presynaptic inhibition than high-threshold mechanoreceptors, thermoreceptors and nociceptors. Cell-type-resolved circuit reconstruction further identified five populations of spinal inhibitory neurons (IhNs) that form synapses onto sensory neuron axons, each preferentially driven by and presynaptically inhibiting a single sensory subtype — a homotypic, modality-specific inhibitory motif. Most of these IhNs also feedforwardly inhibit other sensory pathways — a heterotypic, cross modality motif. Together, dSC1 and SPINE enable cell-type-resolved spinal cord circuit reconstruction and reveal a dual-mode logic of sensory-evoked inhibition that shapes modality-specific signaling at the earliest stage of somatosensory processing.

## Main

Understanding the brain as an embodied system requires determining how sensory stimuli are represented in the central nervous system (CNS). The spinal cord is the first site in the central nervous system where somatosensory information from the skin, muscles, and internal organs is processed (**Figure 1a**)(Abraira and Ginty, 2013; De Groat, 1986; Tuthill and Azim, 2018). More than a dozen physiologically, functionally, and genetically distinct dorsal root ganglion (DRG) sensory neuron subtypes transduce mechanical, thermal, and chemical stimuli into electrical impulses that are conveyed from the body to the spinal cord (**Figure 1b**) (Abraira and Ginty, 2013; Basbaum et al., 2009; Julius, 2013; Lay and Dong, 2020; Lumpkin and Caterina, 2007). Yet, how this diverse repertoire of sensory neurons is wired into spinal cord circuits that mediate spinal reflexes and outputs to the brain remains unclear.

**Figure 1.**
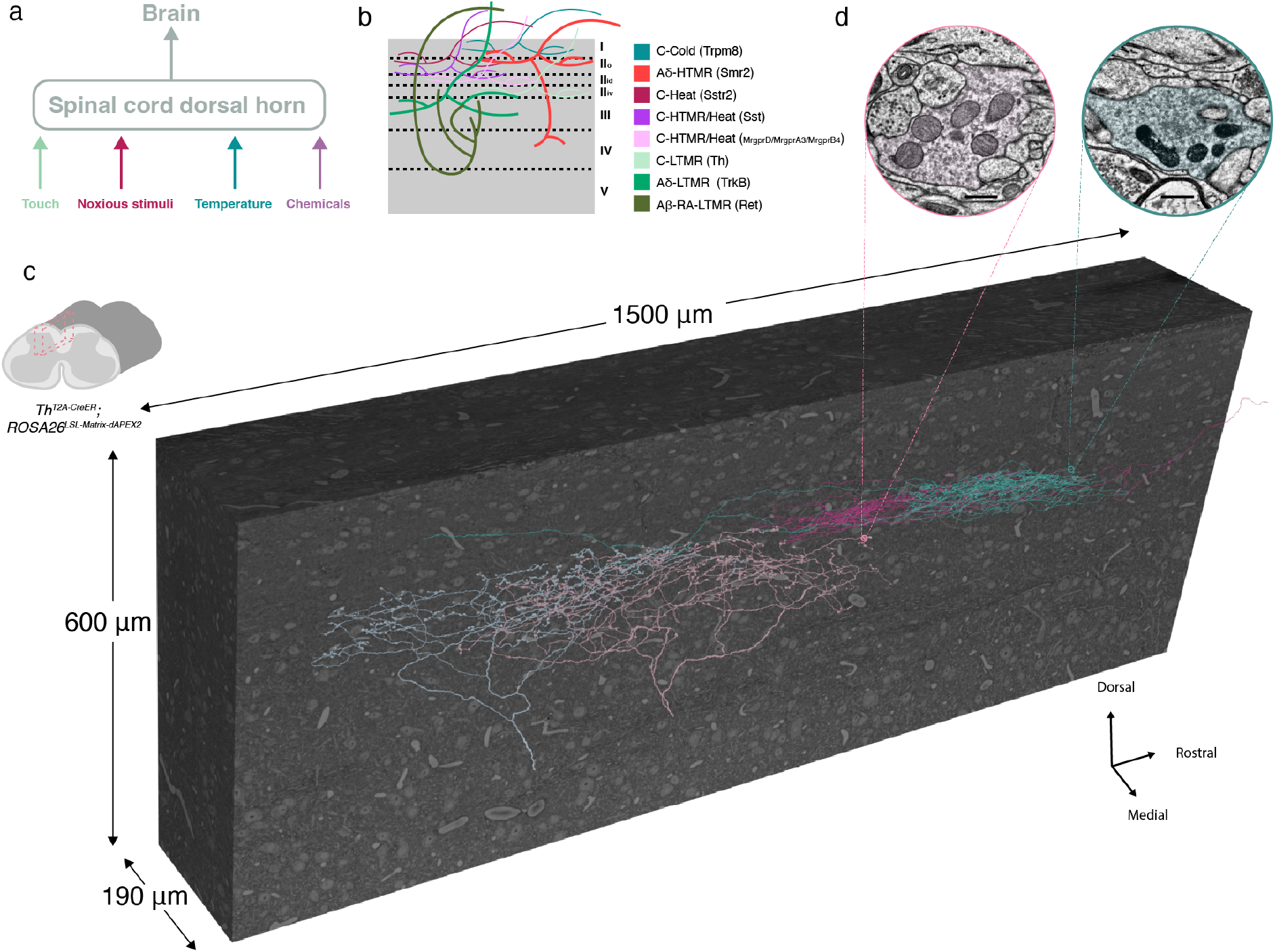
dSC1 EM volumes with genetically labeled sensory neuron subtypes. **a**, Schematic of the somatosensory pathway. The spinal cord dorsal horn receives sensory input from DRG neurons and relays this information to the brain. **b**, Schematic of sensory innervation patterns across laminae of the dorsal horn. **c**, Large-scale EM volume of the lumbar spinal cord dorsal horn. A lumbar spinal cord section was processed from a *Th^T2A-CreER^; ROSA26^LSL-Matrix-dAPEX2^* mouse in which C-LTMRs are labeled by the APEX reporter. The dSC1 EM volume spans 1.5 mm in the rostral-caudal axis, 600 μm in the dorsal-ventral axis and 190 μm in the medial-lateral axis. At the front of the EM volume, two reconstructed Aδ-LTMRs (left) and two APEX-labelled C-LTMRs (right) are shown. **d**, Higher-magnification EM images of an Aδ-LTMR bouton (left) and a C-LTMR bouton containing APEX-labeled mitochondria (right). Scale bar, 500 nm.

Large-scale electron microscopy (EM) provides an opportunity to reconstruct spinal cord circuits at synaptic resolution. However, a major challenge in mammalian EM connectomics is the scalable identification of diverse neuronal cell types, particularly because most neurons are only partially reconstructed in EM volumes (Elabbady et al., 2025; Gamlin et al., 2025; Khajeh and Lee, 2025). Cell-type identification in EM has occasionally been achieved based on characteristic ultrastructural features, including distinctive vesicle and mitochondrial morphologies as well as other subcellular specializations (Elabbady et al., 2025; Guillery, 1969; Ribeiro-da-Silva and Coimbra, 1982; Sager et al., 2026). Indeed, classic EM studies of the spinal cord dorsal horn have suggested that distinct sensory afferent subtypes exhibit characteristic ultrastructural features that may be detectable at the EM level (Rethelyi et al., 1982; Ribeiro-da-Silva and Coimbra, 1982; Ribeiro-da-Silva et al., 1985; Ribeiro-da-Silva et al., 1989; Zhang et al., 2019). Systematically assigning somatosensory afferents in EM datasets to physiologically defined classes based on ultrastructural features could therefore enable cell type-resolved connectomic reconstruction of the synaptic organization and circuit logic of the spinal cord dorsal horn.

Here, we present a large-scale EM volume of the mouse spinal cord dorsal horn, the Spinal Cord dorsal horn 1 (dSC1) dataset, together with a SPinal cord Integrated NEuron classifier (SPINE) trained on the morphological and ultrastructural features of genetically labeled classes of DRG somatosensory neurons. SPINE enabled high-accuracy identification of axons and synapses of eight physiologically distinct DRG sensory neuron subtypes within the dSC1 dataset. Applying SPINE to the dSC1 dataset, we defined the synaptic properties of the different somatosensory neuron subtypes, including presynaptic and feedforward inhibitory motifs with which they associate. Our findings reveal a remarkable heterogeneity of synaptic architecture across the major somatosensory neuron classes and a circuit logic underlying sensory-evoked inhibition for processing somatosensory information within the dorsal horn.

### A large-scale EM volume of mouse lumbar spinal cord dorsal horn

We acquired a serial-section EM volume of the lumbar spinal cord dorsal horn (L3-L4) at synapse-resolution (4 × 4 × 45 nm^3^) from a female *Th^T2A-CreER^; ROSA26^LSL-Matrix-dAPEX2^* mouse, in which C-LTMRs are selectively labeled with a mitochondrial matrix-targeted dAPEX2 reporter (Abraira et al., 2017; Zhang et al., 2019) to allow identification of C-low threshold mechanoreceptor (C-LTMR) axons under EM (**Figure 1d**). Using the GridTape automated sectioning and transmission EM imaging platform (Phelps et al., 2021), we collected and imaged 4,105 sagittal serial-sections of the spinal cord dorsal horn. The resulting EM dataset, named dSC1, spans 1.5 mm along the rostral–caudal axis, 190 µm along the medial–lateral axis, and 600 µm along the dorsal–ventral axis (**Figure 1c**). We then trained convolutional neural networks (CNNs) for automated cell segmentation and reconstruction (Azevedo et al., 2024; Lee et al., 2017). The large size of dSC1, together with automated segmentation, enabled rapid reconstruction of the complete central arbors of both small-caliber (C-fiber) and medium-to-large-caliber (A-fiber) sensory neurons (**Figure 1c**). A single C-LTMR axon can extend up to 6 mm within the EM volume, forming extensively branched arbors. To automatically identify synapses, we trained separate CNNs to detect synapses and assign their presynaptic and postsynaptic compartments (Azevedo et al., 2024; Turner et al., 2022). Automated synapse prediction identified sensory neuron-to-spinal cord neuron synapses with 95.6% precision (the fraction of predicted synapses that were manually confirmed) and 97.3% recall (the fraction of synapses successfully detected), and spinal cord neuron-to-spinal cord neuron synapses with 88.3% precision and 95.5% recall (**Figure S1a**). We found that automated prediction of axoaxonic synapses formed by spinal cord neurons onto sensory neurons achieved high recall but lower precision, likely due to misclassification of some membrane appositions as synapses with certain sensory neuron subtypes (**Figure S1b,c**). We therefore manually validated axoaxonic synapses where necessary. Overall, dSC1, together with automated cell segmentation and synapse prediction, enables rapid, large-scale reconstruction of spinal cord dorsal horn circuits.

### A SPinal cord Integrated NEuron classifier (SPINE)

We next sought to develop a sensory neuron classifier that can be used to identify physiologically distinct sensory neuron subtypes in the dSC1 dataset based on their ultrastructural features. We established a workflow in which genetically labeled sensory neuron subtypes were first used to define subtype-specific ultrastructural features, followed by expert annotation of sensory neuron identities in the dSC1 dataset to generate a training set for classifier development (**Figure 2a**).

**Figure 2.**
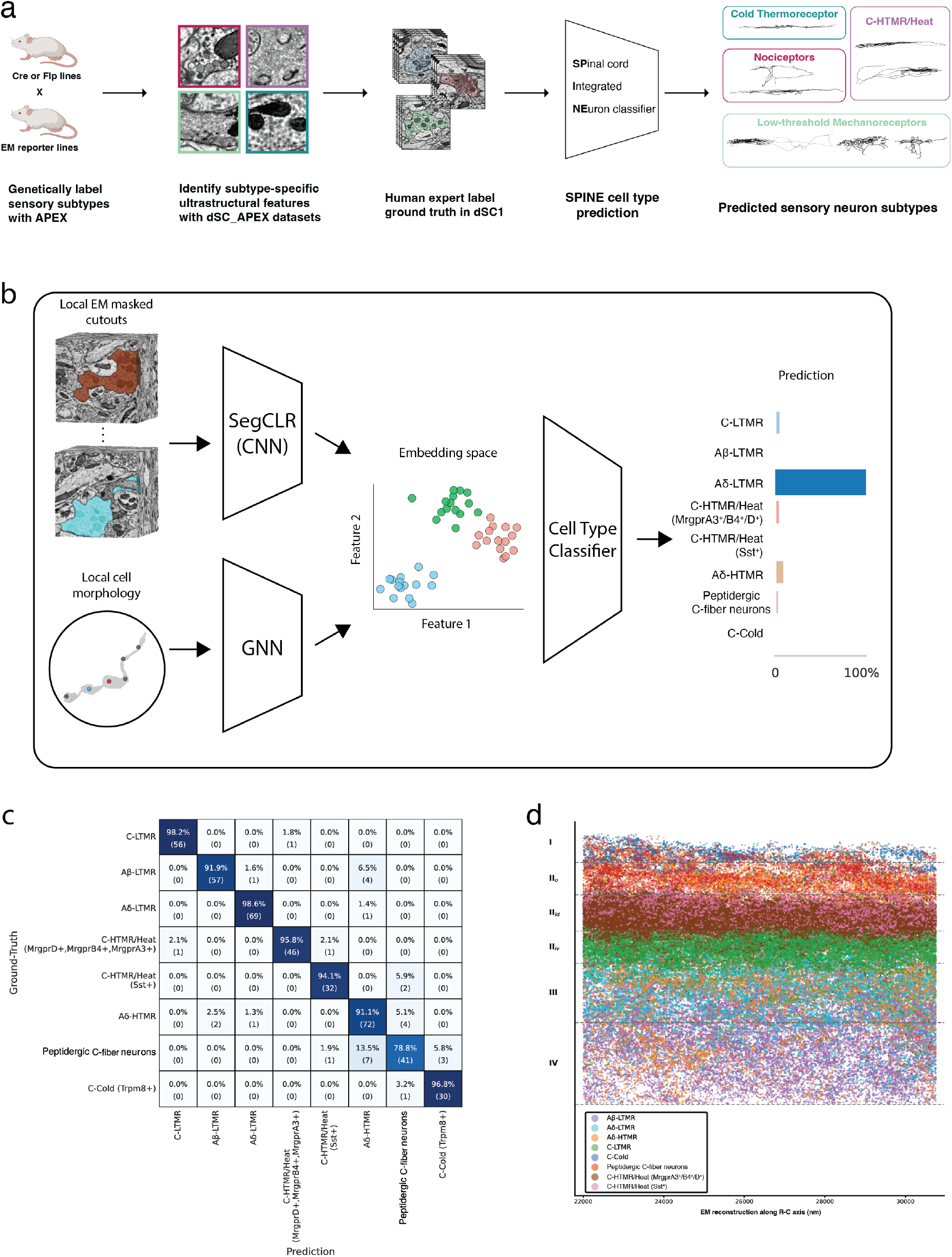
The SPinal cord Integrated NEuron classifier (SPINE) **a**, Machine-learning pipeline to identify major sensory neuron subtypes in EM volumes. Using EM reporters and mouse DRG genetic tools, we generated the dSC_APEX datasets to characterize sensory subtype specific ultrastructural features. We then identified sensory neurons with these features in the dSC1 dataset to generate ground-truth labels and trained a classifier to predict cell-type identity. **b**, Schematic of stage 2 of the SPINE architecture. The framework integrates ultrastructural and morphological features to classify presynaptic sites. (Top left) 3D EM cutouts are processed by SegCLR to generate rich ultrastructural embeddings. (Bottom left) The corresponding synapse mesh is passed through a Graph Neural Network to generate a morphological embedding. (Right) Both embeddings are concatenated and fed through a lightweight MLP for sensory neuron subtype classification. **c**, Confusion matrix of the sensory neuron subtype classifier on a held-out test set. Unnormalized counts of synapses of each category are in parentheses. **d**, Spatial distribution of predicted sensory neuron presynaptic sites across the dorsal horn.

We used DRG neuron genetic tools (Abraira et al., 2017; Han et al., 2013; Lezgiyeva et al., 2026; Li et al., 2011; Luo et al., 2009; Olson et al., 2017; Qi et al., 2024; Rutlin et al., 2014; Sharma et al., 2020; Vrontou et al., 2013) to drive expression of a mitochondria-targeted dAPEX2 EM reporter (Zhang et al., 2019) in select sensory neuron populations in separate datasets. This genetic labeling approach enabled visualization and unambiguous identification of the axons of specific DRG neuron populations within the EM volumes. Thus, we generated six additional large-scale EM datasets, each labeling one or two genetically defined sensory neuron subtypes, collectively termed the dSC_APEX datasets (**Figure S2a,b**). Using the six dSC_APEX datasets, together with single-section, APEX-labeled TEM datasets to visualize axons and synapses of MrgprB4⁺ and Sst⁺ C-high threshold mechanoreceptor (HTMR)/Heat neurons, we characterized ultrastructural features across ten genetically labeled DRG sensory neuron subtypes. The ten physiologically distinct and genetically labeled sensory neuron subtypes were: C-fiber cold thermoreceptors (TrpM8^+^ C-Cold); C-fiber heat thermoreceptors (Sstr2^+^; CGRP^+^ C-Heat nociceptors); Aδ-HTMRs (Smr2^+^; CGRP^+^ Aδ-HTMR or myelinated nociceptors); C-HTMR/Heat (Sst^+^) neurons; C-HTMR/Heat (MrgprD^+^) neurons; C-HTMR/Heat (MrgprA3^+^) neurons; C-HTMR/Heat (MrgprB4^+^) neurons; C-LTMRs; Aδ-LTMRs; and Aβ RA-LTMRs. Together, these APEX labeling experiments allowed us to characterize the ultrastructural features of neurons that represent more than 90% of DRG neurons innervating the lumbar spinal cord dorsal horn, excluding proprioceptors (estimated from (Sharma et al., 2020)). We reconstructed the labeled axon terminals in the volumetric dSC_APEX datasets using automated segmentation to visualize their terminal morphologies.

This analysis revealed that sensory neuron subtypes are distinguishable based on synaptic vesicle morphology, bouton shape, myelination pattern, and laminar location (**Figures S3, S4; Methods**). Trpm8⁺ C-cold thermoreceptors terminated in the dorsal-most region of lamina I and contained predominantly round, clear synaptic vesicles, suggesting they contain mostly glutamatergic vesicles (**Figure S3a**) (Peters and Palay, 1996). Ventral to these, peptidergic (CGRP^+^) C and Aδ-fibers terminated primarily in lamina I and outer lamina II and exhibited vesicle populations distinct from the small round clear vesicles found in canonical glutamatergic terminals. Sstr2⁺ C-heat nociceptors contained abundant dense-core vesicles (∼100 nm diameter) (**Figure S3b)**, whereas myelinated Smr2⁺ Aδ-HTMR nociceptors contained characteristic large pale vesicles (∼200 nm diameter) within their axonal boutons (**Figure S3c)** and additionally gave rise to collaterals that terminated in laminae III and IV, as recently observed by light microscopy (Lezgiyeva et al., 2026). Among the four C-HTMR/Heat subtypes, which terminated in lamina II inner-dorsal (IIid), Sst⁺ C-HTMR/Heat neurons were uniquely distinguished by elongated spindle-shaped vesicles (**Figure S3d)**, whereas MrgprA3⁺, MrgprD⁺, and MrgprB4⁺ C-HTMR/Heat neurons shared ultrastructural features of the classic type I glomerulus, including dense clusters of variably sized, pale vesicles (Ribeiro-da-Silva and Coimbra, 1982; Zhang et al., 2019) (**Figure S3e,f)**.

LTMRs terminated ventral to lamina IIid and, like Trpm8⁺ C-cold thermoreceptors, contained predominantly round, clear vesicles (**Figure S3g-i)**. However, C-LTMRs, Aδ-LTMRs, and Aβ RA-LTMRs could be distinguished by their neurofilament abundance (**Figure S3g-i; Figure S4g,h)**, terminal morphology (**Figure S4d-f**), and myelination patterns (**Figure S4g.h)**. Three-dimensional reconstructions further revealed distinct subtype-specific bouton architectures. For example, MrgprA3⁺, MrgprD⁺, and MrgprB4⁺ C-HTMR/Heat afferents frequently formed perforated boutons that were deeply invaginated by pre- and postsynaptic partners (**Figure S4c**), whereas C-LTMRs and Aδ-LTMRs exhibited a characteristic "beads-on-a-string" morphology with boutons connected by thin inter-bouton segments (**Figure S4d,f**). In contrast, Aβ RA-LTMR terminals were larger in diameter and formed fewer discrete boutons (**Figure S4e**).

These ultrastructural features were used by expert annotators to identify sensory neuron subtypes directly within the large dSC1 dataset, generating a ground-truth training set for model training and validation. Rather than transferring annotations from the dSC_APEX datasets, we established training labels directly in dSC1 because independently acquired EM datasets can differ in intrinsic imaging characteristics and, importantly, mitochondrial APEX labeling in the dSC_APEX datasets introduced dataset-specific ultrastructural features that could bias classifier training (**Figure 2a**). Instead, the dSC_APEX datasets were used as independent validation benchmarks to test how well the classifier generalized across different animals and experimental conditions.

Our ground-truth dataset in dSC1 comprised 2,057 sensory neuron cutouts (local fragments computationally extracted from segmented neurons), each containing the corresponding EM images and 3D mesh and centered on a presynaptic site (**Methods**). Each presynaptic sensory neuron cutout was assigned to one of eight sensory neuron classes based on ultrastructural features: C-Cold thermoreceptors; peptidergic (CGRP^+^) C-fiber neurons (peptidergic C-fiber neurons); Aδ-HTMRs (myelinated mechano-nociceptors); Sst⁺ C-HTMR/Heat neurons; MrgprA3⁺/MrgprD⁺/MrgprB4⁺ C-HTMR/Heat neurons; C-LTMRs; Aδ-LTMRs; and Aβ-LTMRs. We combined MrgprA3⁺, MrgprD⁺, and MrgprB4⁺ C-HTMR/Heat neurons into a single class because they could not be reliably distinguished at the EM level without APEX labeling. We also could not distinguish the Aβ-LTMR subtypes (Aβ SA-LTMRs, Aβ RA1-LTMRs, Aβ RA2-LTMRs, and Aβ field-LTMRs) from one another due to limitations in available genetic tools. Therefore, large-diameter myelinated LTMRs with abundant neurofilament and morphologies resembling APEX-labeled Aβ RA-LTMR terminals were classified collectively as Aβ-LTMRs. Similarly, we could not resolve peptidergic C-fiber neuron subtypes (C-thermo-nociceptors, labeled using *Sstr2^CreER^* mice; Adra2a^+^; and Oprk1^+^ peptidergic C fiber neuron subtypes (Qi et al., 2024; Sharma et al., 2020)). Thus, unmyelinated C-fibers terminating in lamina I and outer lamina II that contained dense-core vesicles and resembled APEX-labeled Sstr2⁺ C-Heat nociceptor terminals were classified as peptidergic C-fiber neurons. Moreover, proprioceptors were excluded because their central terminals lie in more ventral regions of the spinal cord. Despite these limitations, the ground-truth dataset encompasses the major DRG sensory neuron classes.

Using this ground truth, we developed the SPinal cord Integrated NEuron classifier (SPINE), a deep learning-based classifier that predicts sensory neuron subtypes based on their ultrastructural and morphological features, described above (**Figure 2b**). Instead of requiring fully proofread or completely reconstructed sensory axon arbors, SPINE operates on local presynaptic fragments by using masked 3D EM cutouts as input. SPINE operates in two sequential stages. First, a lightweight binary classifier uses segmentation-guided contrastive learning of representations (SegCLR) (Dorkenwald et al., 2023) to discriminate between presynaptic neuronal fragments belonging to sensory neurons versus non-sensory neurons (**Figure S5a**). Second, for presynaptic fragments classified as sensory, SPINE expands its spatial window to a 5-10 µm radius to capture broader synapse features. To predict specific sensory neuron subtypes, this stage extracts dual-contrastive ultrastructural features using SegCLR models trained on both the MICrONS (Consortium, 2025) and H01 (Shapson-Coe et al., 2024) datasets and concurrently processes the local synapse mesh geometry through a Graph Neural Network (GNN) (Kipf and Welling, 2017). These ultrastructural and morphological features were concatenated into a unified vector to assign each segment to one of eight sensory neuron subtypes. SPINE generated a probability for each class, and predictions with low maximum probability were labeled as low confidence, which likely captures sensory afferent subtypes not represented in the training set (usually fewer than 5%, **Figure S5b**). Following training, the sensory neuron subtype classifier achieved 93% overall accuracy, with per-class accuracy ranging from 91–99%, except for peptidergic C-fiber neurons (78%; **Figure 2c**). When applied to randomly sampled presynaptic sensory neuron segments across the dorsal horn in dSC1, the inferred laminar distributions of the sensory neuron subtypes closely matched those defined in previous light microscopy studies (**Figure 2d**) (Abraira et al., 2017; Lezgiyeva et al., 2026; Qi et al., 2024).

To further test whether SPINE trained on dSC1 ground truth data indeed learned to identify the correct sensory neuron subtypes in EM volumes, we evaluated the accuracy of the model on the dSC_APEX datasets in which different sensory neuron subtypes were APEX-labeled and segmented (**Figure S5c,d**). SPINE predicted cell types in these datasets with more than 70% accuracy, even though the dSC_APEX datasets were collected from different animals and contained APEX-labeled mitochondria that were absent from the ground-truth dataset. These results confirm that SPINE learned to accurately identify each of the eight major sensory neuron subtypes in EM volumes and can be used beyond the dSC1 dataset.

To better understand the local features SPINE uses for its predictions, we applied Quantitative Attribution with Counterfactuals (QuAC) (Adjavon et al., 2024). QuAC generates visual explanations by minimally modifying an image to shift a classifier’s prediction to an alternative class, thereby identifying class-relevant differences. Because QuAC was designed for 2D-based models, we trained a specialized 2D ResNet-18 classifier that predicts sensory neuron subtypes from a single serial-section EM image centered on the presynaptic sites (**Figure S5e**). We then used QuAC to convert one sensory neuron subtype into another.

We found that QuAC primarily altered the characteristic vesicle morphologies of sensory neurons to change the model’s prediction. For example, when converting an Aδ-HTMR into a peptidergic C-fiber neuron, QuAC transformed large pale vesicles into dense-core vesicles. Conversely, when converting a peptidergic C-fiber neuron into an Aδ-HTMR, it replaced dense-core vesicles with large pale vesicles (**Figure S5f,g**). These results align with the subtype-specific ultrastructural signatures observed in the six dSC_APEX datasets and suggest that vesicle morphology within local presynaptic segments is a key feature used by the SPINE classifier to distinguish sensory neuron subtypes.

### Synapse organization across physiologically distinct sensory neuron subtypes

Somatosensory afferents elaborate extensive axonal arbors within the spinal cord (Abraira and Ginty, 2013; Brown, 2012; Kuehn et al., 2019) and form specialized synaptic architectures with dorsal horn neurons (Rethelyi et al., 1982). Pioneering EM studies described glomerular structures formed by sensory afferents. These structures comprise axodendritic synapses, formed by sensory axons onto dorsal horn neurons, and axoaxonic synapses formed by inhibitory spinal interneurons (IhNs) onto sensory neuron axons, which mediate presynaptic inhibition (PSI) (**Figure 3a,b**) (Cruz et al., 1991; Ribeiro-da-Silva and Coimbra, 1982; Ribeiro-da-Silva et al., 1985; Todd, 1996; Todd and Lochhead, 1990). Such a synaptic architecture creates a tightly packed microcomplex of interconnected neurons that serves as a primary sensory processing hub, where the relay of sensory signals to spinal cord neurons is regulated by mechanisms such as PSI (**Figure 3c**). Yet, it remains unclear how this organization may vary across sensory neuron subtypes that convey distinct somatosensory modalities. Therefore, we used dSC1 and SPINE to investigate the distribution and organization of axodendritic and axoaxonic synapses across the physiologically distinct DRG sensory neuron subtypes.

**Figure 3.**
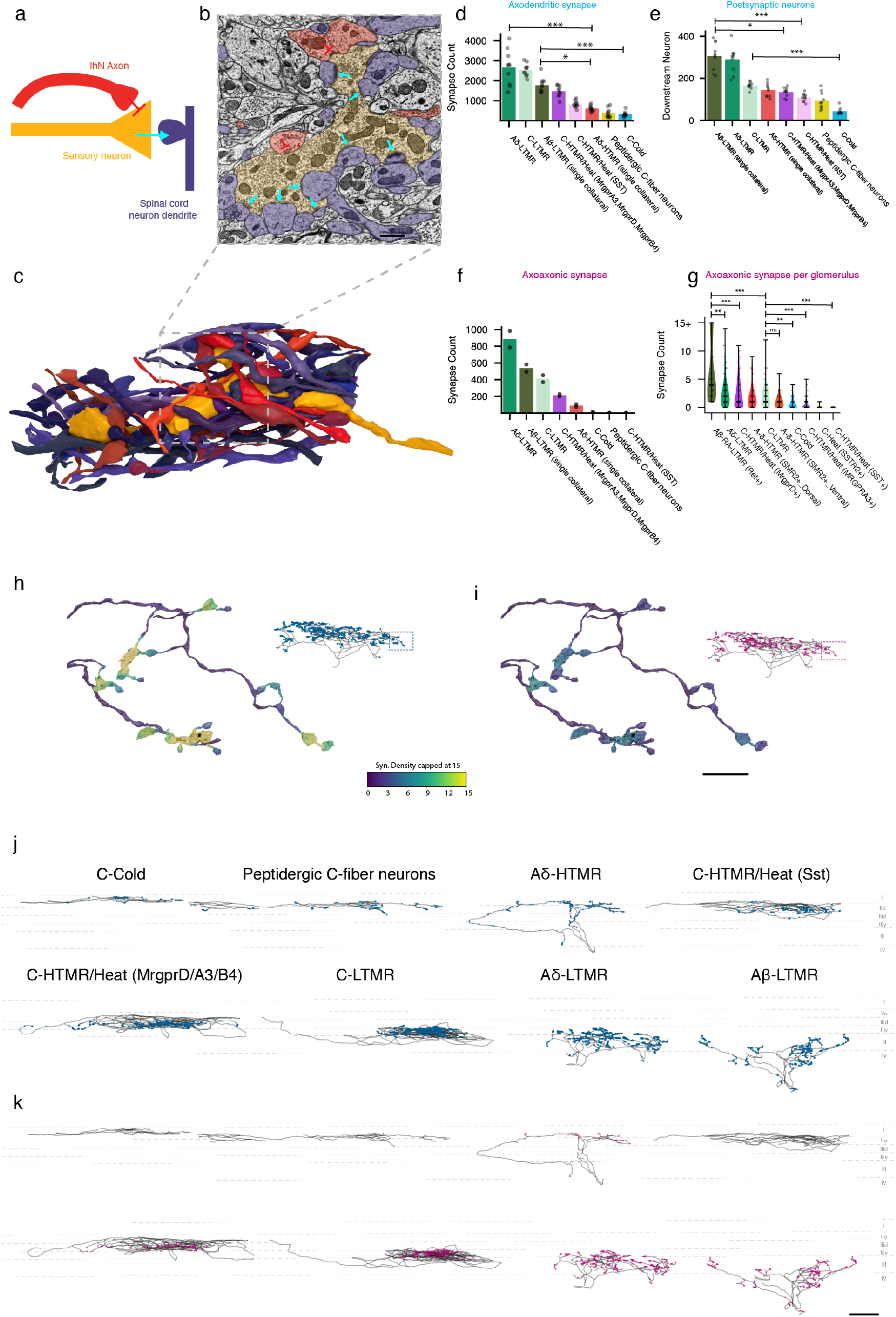
Synaptic architectures of somatosensory neuron subtypes. **a,** Schematics of synapse arrangements of sensory axonal boutons. Sensory neurons make axodendritic synapses onto spinal cord neurons (blue arrow). GABAergic spinal neurons’ axons (red) form axoaxonic synapses onto sensory axons to mediate PSI. **b,** EM images showing an Aδ-LTMR bouton forming axodendritic synapses (blue arrow) and axoaxonic synapses (red arrows). Scale bar, 750 nm. **c**, 3D reconstruction of the complex synaptic organization around the same Aδ-LTMR bouton (yellow) shown in **b**, including 11 presynaptic axons (red) forming axoaxonic synapses onto the Aδ-LTMR bouton and 15 spinal cord neuron dendrites (purple) receiving Aδ-LTMR input via axodendritic synapses. Neuron meshes are cropped to a 4-µm radius around the Aδ-LTMR bouton. **d,** Total axodendritic synapse counts postsynaptic partner neurons for the full central projections of each sensory subtype in the dSC1 dataset were quantified. For Aβ-LTMRs and Aδ-HTMRs, single collaterals were reconstructed and quantified. (Aδ-LTMR versus Aδ-HTMR single collateral, p=4.07 × 10^-4^; Aβ-LTMR collaterals versus C-Cold, p=8.03 × 10^-4^; Aβ-LTMR collaterals versus Aδ-HTMR single collateral, p=4.52 × 10^-2^, Kruskal-Wallis test, Dunn post hoc test. n = 10 neurons per subtype, except C-Cold (n = 9)). **e**, Total number of downstream neurons across sensory neuron subtypes in the dSC1 dataset. (Aβ-LTMR collaterals versus C-HTMR/Heat (Sst), p=2.55 × 10^-4^; Aβ-LTMR collaterals versus C-HTMR/Heat (MrgprD/A3/B4), p=1.52 × 10^-2^; C-LTMR versus C-Cold (Trpm8^+^), p=6.30 × 10^-4^, Kruskal-Wallis test, Dunn post hoc test. n = 10 neurons per subtype, except C-Cold (n = 9)) **f,** Total axoaxonic synapse counts across the full central projections of each sensory subtype in the dSC1 dataset. Axoaxonic synapses were automatically predicted followed by manual validation. **g,** Total axoaxonic synapse counts per single glomerulus in SC_APEX datasets (**Methods**). (Aβ-RA LTMR versus C-LTMR, p=2.74 × 10^-5^; Aβ-RA LTMR versus MrgprD^+^ C-HTMR/Heat, p=1.07× 10^-4^; Aβ-RA LTMR versus Aδ-LTMR, p=5.43 × 10^-3^; C-LTMR versus Sst^+^ C-HTMR/Heat, p=2.75 × 10^-10^; C-LTMR versus MrgprA3^+^ C-HTMR/Heat, p=1.04 × 10^-7^; C-LTMR versus C-Cold, p=2.60 × 10^-3^; C-LTMR versus Aδ-ΗΤΜR (Ventral), p=0.51. Kruskal-Wallis test, Dunn post hoc test. *n* = 42 Aβ-RA-LTMR, 102 Aδ-LTMR, 45 MrgprD^+^ C-HTMR/Heat, 51 dorsal Aδ-HTMR, 79 C-LTMR, 45 ventral Aδ-HTMR, 42 C-Cold, 118 MrgprA3^+^ C-HTMR/Heat, 34 Sstr2^+^ C-Heat and 44 Sst^+^ C-HTMR/Heat glomeruli.). **h**. Axodendritic synapses localize to axon terminals. Left, an Aδ-LTMR axon terminal mesh color-coded by synapse density, measured as the number of axodendritic synapses within a 4 μm radius. Right, skeleton trace of the Aδ-LTMR with axodendritic synapse locations shown as blue dots. Boxes indicate the axon terminals shown on the left. **i**. Same neuron as in **g**, with axoaxonic synapse density and localization shown. (scale bar 10 µm). **j,** Skeleton traces of sensory neuron subtypes with axodendritic synapse locations (blue dots). **k,** Same neuron as in **g**, with individual axoaxonic synapses shown as dots. (scale bar 100 µm).

We first examined the axodendritic synapses through which sensory neurons relay information to spinal cord interneurons and projection neurons. We reconstructed the central arbors of each of the eight sensory neuron subtypes in the dSC1 dataset. For Aβ-LTMRs and Aδ-HTMRs, which branch when they enter the spinal cord and extend along the rostral-caudal axis to form multiple gray matter penetrating axon collaterals (Abraira and Ginty, 2013; Bai et al., 2015; Brown et al., 1977, 1978; Kuehn et al., 2019; Lezgiyeva et al., 2026), we reconstructed individual penetrating axon collaterals. The number of axodendritic synapses varied markedly across sensory neuron subtypes. LTMRs, as a population, formed significantly more axodendritic synapses than Aδ-HTMRs, peptidergic C-fiber neurons, and C-cold thermoreceptors (**Figure 3d,j**). On average, individual Aδ-LTMRs and C-LTMRs formed 2,662 ± 86 and 2,532 ± 316 axodendritic synapses, respectively. A single Aβ-LTMR collateral formed an average of 1,772 ± 360 axodendritic synapses. In fact, since Aβ-LTMRs have 10 or more penetrating axon collaterals (Bai et al., 2015; Brown et al., 1980; Kuehn et al., 2019; Li et al., 2011), individual Aβ-LTMRs have more than 17,000 axodendritic synapses. By contrast, individual C-cold thermoreceptors formed only 320 ± 138 axodendritic synapses. The number of postsynaptic partner neurons also varied across sensory neuron subtypes: Individual Aβ-LTMR collaterals and Aδ-LTMR neurons contacted on average ∼300 spinal cord neurons (i.e. ∼3,000 or more postsynaptic partner neurons of individual Aβ-LTMRs because of their 10 or more collaterals), which is considerably more than individual C-HTMR/Heat neurons, peptidergic C-fiber neurons, and C-Cold neurons (**Figure 3e**). Thus, sensory neuron subtypes exhibit markedly different numbers of synapses and postsynaptic partner neurons, with A-fiber LTMRs forming the most synapses and broadcasting their signals onto the largest number of postsynaptic partners.

We next investigated the number and distribution of axoaxonic synapses along the axons of reconstructed sensory neuron subtypes. The three LTMR classes exhibited the highest number of axoaxonic synapses, with a single C-LTMR, Aδ-LTMR, and Aβ-LTMR axon collateral typically receiving >400 axoaxonic synapses. Conversely, Sst⁺ C-HTMR/Heat, peptidergic C-fiber neurons, and C-cold neurons were virtually devoid of axoaxonic synapses (**Figure 3f,k**). Because the SPINE classifier does not distinguish MrgprA3⁺, MrgprD⁺, and MrgprB4⁺ C-HTMR/Heat neurons or specific peptidergic C-fiber neuron subclasses in the dSC1 dataset, we also analyzed the dSC_APEX volumes in which these populations were independently labeled by APEX (**Figure 3g**). We approximate glomerular structures by clustering synapses in close proximity (**Methods**) and then quantified the number of axoaxonic synapses received by individual sensory glomeruli. Consistent with the whole-axon analysis, Aβ RA-LTMR glomeruli received the greatest axoaxonic innervation, with a median of 4 axoaxonic synapses per glomerulus (IQR 2-9). In comparison, Aδ-LTMR, C-LTMR, MrgprD⁺ C-HTMR/Heat neuron, and dorsal Aδ-HTMR glomeruli received intermediate levels of axoaxonic innervation (median 1-2 synapses, IQR 0-4), whereas C-cold, MrgprA3⁺ C-HTMR/Heat, Sstr2⁺ C-heat, and Sst⁺ C-HTMR/Heat neuron glomeruli, and ventral axonal branches of Aδ-HTMRs received few or no axoaxonic synapses (median 0-1 synapse, IQR 0-1). Together, these analyses demonstrate that LTMRs receive many axoaxonic synapses to govern GABA-mediated PSI, whereas MrgprA3⁺ and Sst⁺ C-HTMR/Heat neurons and two thermoreceptor classes (C-heat and C-cold) receive few to no axoaxonic synapses and are thus minimally modulated by GABA-mediated PSI.

We next investigated how axodendritic and axoaxonic synapses are distributed along the axonal arbors of sensory neuron subtypes. Across all sensory neuron subtypes, axodendritic synapses— and, for sensory neuron subtypes that receive PSI, axoaxonic synapses—were highly enriched at distal axon terminals rather than being uniformly distributed along their axonal arbors (**Figure 3h,i; Figure S6a**). For all three LTMR subtypes, ∼75% of both axodendritic and axoaxonic synapses were located within 25 μm of an axonal tip, despite this distal region comprising only ∼20% of their total axonal arbor in the dorsal horn gray matter (**Figure S6b,c**). Aδ-HTMRs, peptidergic C-fiber neurons, and C-cold thermoreceptors formed more *en passant* axodendritic synapses, with ∼75% of synapses located within 50 μm of an axonal tip.

The strong enrichment of both axodendritic and axoaxonic synapses within confined distal terminal regions resulted in extensive colocalization of these two synapse types. In LTMRs, virtually all axodendritic synapses were located within 5 μm of an axoaxonic synapse (**Figure S6d-h**). As a result, individual axodendritic synapses of Aβ-LTMRs were accompanied by, on average, >9 axoaxonic synapses within a 4 μm distance. Aδ-LTMRs, C-LTMRs, and MrgprD⁺ C-HTMR/Heat neurons exhibited a similar organization, with typically >3 axoaxonic synapses located within 4 μm of an axodendritic synapse (**Figure S6i**). Interestingly, Smr2⁺ Aδ-HTMR axons, which extend and form synapses across laminae I–IV, exhibited lamina-specific differences in the distribution of axoaxonic synapses and thus GABA-mediated PSI. Dorsal terminal branches contained more axoaxonic synapses in proximity to axodendritic synapses than ventral branches of the same axon (**Figure S6i,j**), suggesting that PSI of Aδ-HTMRs is differentially regulated across laminar targets.

These findings indicate that among sensory neuron subtypes that receive axoaxonic synapses, these synapses colocalize extensively with axodendritic synapses at distal afferent terminals. This spatial coupling of axodendritic and axoaxonic synapses provides an anatomical substrate for GABA-mediated PSI to modulate transmitter release at the majority of release sites on a sensory neuron. Moreover, the abundance of axoaxonic synapses that form upon all three LTMR classes compared to the near absence of axoaxonic synapses on thermoreceptors (Trpm8⁺ C-cold and Sstr2⁺ C-heat) and on itch-associated C-HTMR/Heat populations (MrgprA3⁺ and Sst⁺) is striking and reveals that GABA-mediated presynaptic modulation is deployed differently across the sensory modalities.

### Physiological validation of subtype differences in PAD

We next used slice electrophysiology as a test of predictions made by the sensory neuron subtype axoaxonic synapse findings across the sensory neuron subtypes in the EM datasets. In the dorsal horn, interneurons that form axoaxonic synapses onto sensory neuron terminals release GABA, which activates ionotropic GABA_A_ receptors and depolarizes the sensory axons (Eccles et al., 1962; Zimmerman et al., 2019); this process is termed primary afferent depolarization (PAD). We expressed channelrhodopsin-2 in dorsal horn GABAergic interneurons using Vgat-ChR2 mice and optogenetically activated these neurons while recording PAD-evoked back-propagating action potentials to the somata of genetically labeled sensory neuron subtypes (**Figure S7a**). We observed that broad activation of spinal cord GABAergic neurons reliably evoked back-propagating action potentials in Aβ RA-LTMRs, Aδ-LTMRs, and C-LTMRs (**Figure S7b-e, j**).

Consistent with these responses being mediated by GABA, application of a GABA_A_ receptor antagonist eliminated the light-evoked back-propagating action potentials (**Figure S7h,i**). In contrast, CGRP⁺ peptidergic C-fiber neurons failed to fire back-propagating action potentials upon GABAergic neuron activation (**Figure S7f,g**). These findings are consistent with our reconstructions showing that LTMRs receive abundant axoaxonic synapses whereas peptidergic C-fiber neurons largely lack them. Interestingly, we failed to evoke action potentials in MrgprD⁺ C-HTMR/Heat neurons and CGRP⁺ Aδ-HTMRs by activating GABAergic neurons (**Figure S7j**). This likely reflects the relatively sparse axoaxonic synapses associated with these populations, compared with LTMRs (**Figure 3c**). Consistent with this interpretation, when we expressed the light-gated chloride channel Acr1 in sensory neurons and used light to directly increase chloride conductance in their central terminals, optogenetic activation reliably evoked back-propagating action potentials in A- and C-fibers, including C-HTMRs (**Figure S7k-n**), indicating that these afferents are indeed capable of generating PAD when sufficient chloride current is provided. These findings from electrophysiological experiments corroborate our EM reconstruction observations that LTMRs receive far more axoaxonic synapses and thus exhibit stronger GABA-mediated PSI compared to other somatosensory neuron subtypes.

### Homotypic presynaptic inhibitory circuits

Our structural and physiological characterization of PSI mediated by GABAergic axoaxonic synapses highlights the central role of spinal inhibitory neurons (IhNs) in shaping sensory processing in a modality-specific manner (Boyle et al., 2019; Chirila et al., 2022; Davis et al., 2023; Foster et al., 2015; Kardon et al., 2014; Melzack and Wall, 1965; Ross et al., 2010; Russ et al., 2021; Zhang et al., 2025; Zimmerman et al., 2019). At the circuit level, dorsal horn IhNs that mediate PSI (PSI-IhN) by forming axoaxonic synapses onto sensory afferent terminals (Todd, 1996), as characterized above, also provide feedforward inhibition (FFI) of spinal cord neurons through axodendritic or dendrodendritic synapses (**Figure 4a**) (Boyle et al., 2019; Chirila et al., 2022; Hughes et al., 2012). Therefore, we next sought to identify PSI-IhNs in the dSC1 dataset and define how their synaptic connections are organized with respect to the sensory neuron subtypes they engage.

**Figure 4.**
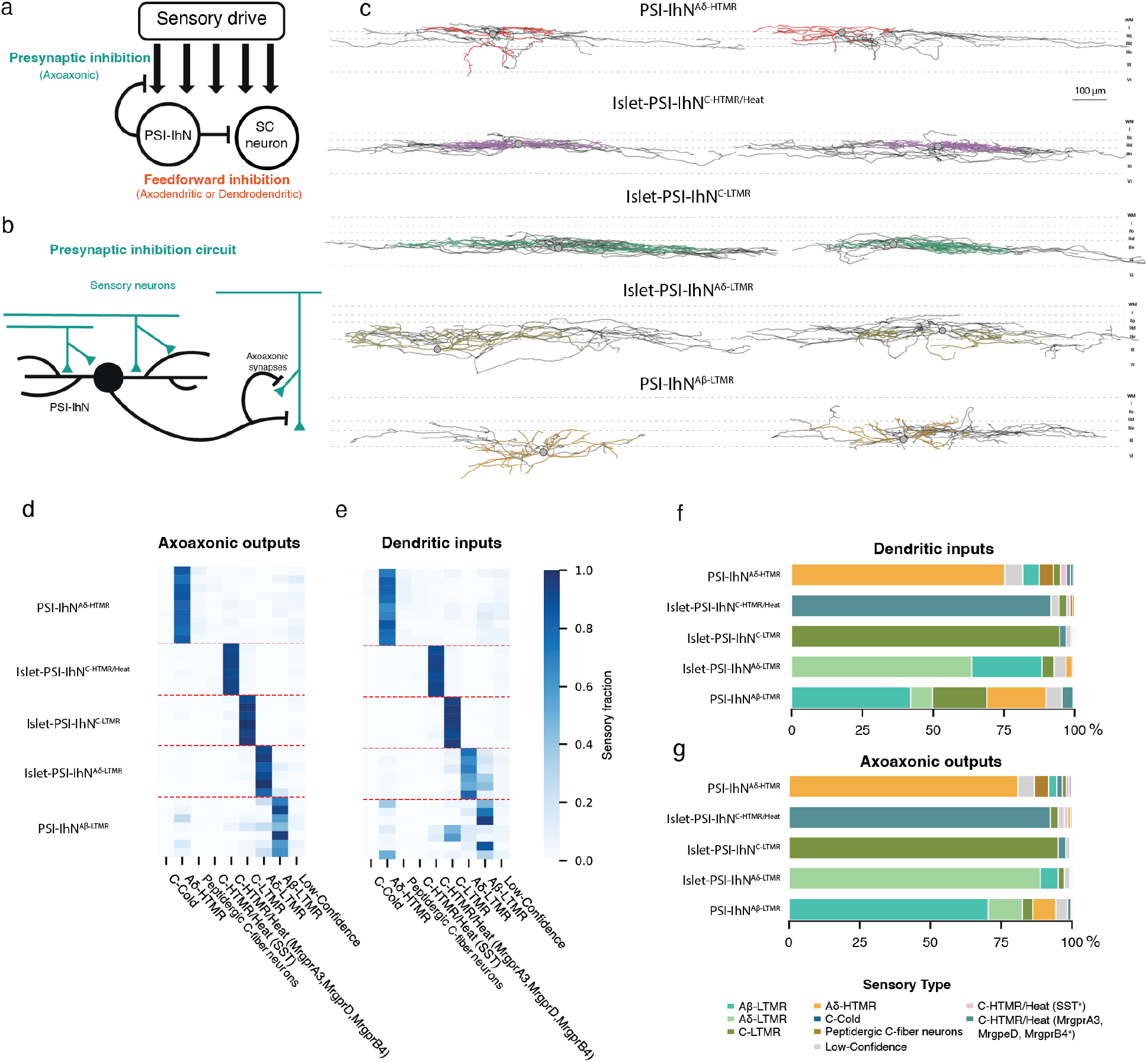
Spinal circuit motifs mediating sensory-evoked homotypic presynaptic inhibition (PSI) across sensory neuron subtypes. **a,** Schematic of sensory evoked PSI and FFI. **b**, Schematic of homotypic PSI circuits. Inhibitory dorsal horn neurons receive input from a given sensory subtype and form axoaxonic synapses back onto axons of the same subtype, thereby mediating PSI. **c**, Skeleton reconstructions of five PSI-IhN neurons. Dendrites are shown as colored skeletons, axons as black skeletons, and somata as gray circles. **d**, Fractions of outputs from each inhibitory subtype onto different sensory subtypes. **e**, Fractions of inputs from each sensory subtype onto inhibitory neuron subtypes (rows) that mediate PSI. **f,** Summed sensory input fractions onto each PSI-IhN population (from **e**). **g,** Summed axonal output fractions from each PSI-IhN population (from **d**).

We identified and reconstructed 34 of the PSI-IhNs across lamina I-III that form axoaxonic synapses onto axons of Aβ-LTMRs, Aδ-LTMRs, C-LTMRs, Aδ-HTMRs, and C-HTMR/Heat neurons (**Figure 4c**). Remarkably, the axons of these PSI-IhNs preferentially formed synapses upon a single sensory neuron subtype: in most cases, more than 70% of their axoaxonic synapses were made onto axons of one sensory neuron class (**Figure 4d; Figure S8a**). PSI-IhNs forming axoaxonic synapses upon C-HTMR/Heat neuron, C-LTMR, and Aδ-LTMR axons formed a morphologically coherent group of neurons with elongated dendritic arbors, which are the previously described islet cells (**Figure 4c**) (Davis et al., 2023; Grudt and Perl, 2002; Todd and Lewis, 1986). In contrast, PSI-IhNs forming axoaxonic synapses onto Aβ-LTMRs and Aδ-HTMRs exhibited more heterogeneous dendritic morphologies. Based on their morphologies and axoaxonic output selectivity, we designated these GABAergic interneuron populations Islet-PSI-IhN^C-HTMR/Heat^, Islet-PSI-InN^C-LTMR^, Islet-PSI-IhN^Aδ-LTMR^, PSI-IhN^Aδ-HTMR^, and PSI-IhN^Aβ-LTMR^.

We next applied the SPINE model to identify the presynaptic sensory neuron synaptic inputs to each of these five PSI-IhN populations and aligned these inputs with their axoaxonic output partners. This analysis revealed a remarkable and generalizable circuit motif: PSI-IhNs that formed axoaxonic synapses received most of their excitatory input from the same sensory subtype that they inhibit via PSI (**Figure 4d-g; Figure S8b**). For example, Islet-PSI-IhN^C-HTMR/Heat^ neurons received >90% of their sensory input from C-HTMR/Heat afferents and formed the vast majority of their axoaxonic synapses onto C-HTMR/Heat terminals. Similar homotypic motifs were observed for all Islet-PSI-IhN^C-LTMR^, Islet-PSI-IhN^Aδ-LTMR^, PSI-IhN^Aδ-HTMR^, and a subset of PSI-IhN^Aβ-LTMR^ neurons. We refer to this form of sensory neuron subtype-specific inhibitory feedback as homotypic PSI (Zimmerman *et al*., 2019).

Notably, although PSI-IhN^Aβ-LTMR^ neurons directed most of their axoaxonic output to Aβ-LTMRs, their sensory inputs were more heterogeneous than those of other PSI-IhN populations. Three out of seven PSI-IhN^Aβ-LTMR^ neurons received predominantly Aβ-LTMR input, consistent with homotypic PSI. However, C-LTMRs provided the largest sensory input to two PSI-IhN^Aβ-LTMR^ neurons, accounting for 58.2% and 46.8% of their total sensory input, while these neurons still preferentially formed axoaxonic synapses onto Aβ-LTMRs. Two additional PSI-IhN^Aβ-LTMR^ neurons received substantial Aδ-HTMR input, accounting for 37.4% and 49.5% of their sensory input.(**Figure 4d,e**). Thus, in addition to homotypic PSI, these circuits provide a potential route through which C-LTMRs and fast-conducting nociceptors can presynaptically gate Aβ-LTMR input through local spinal GABAergic interneurons. Moreover, beyond primary afferent drive, these PSI-IhNs also received substantial input from spinal interneurons and other CNS sources (Abraira *et al*., 2017). Interestingly, PSI-IhN^Aδ-HTMR^ and PSI-IhN^Aβ-LTMR^ neurons exhibited significantly less direct sensory input than the islet-like populations (**Figure S8c**), suggesting that PSI of Aδ-HTMR and Aβ-LTMR terminals is more strongly controlled by central spinal cord circuit computations.

Although the homotypic inhibitory motifs described above suggest subtype-selective PSI at the population level, PSI could in principle arise from feedback onto an afferent’s own terminals, from lateral inhibition of different afferent terminals of the same subtype, or from a combination of both (**Figure S9a**). To distinguish between these possibilities, we quantified the extent of connectivity between individually reconstructed sensory neurons and individually reconstructed PSI-IhNs by calculating, for each sensory neuron–PSI-IhN pair, the fraction of the PSI-IhN’s sensory input contributed by that sensory neuron and the fraction of its axoaxonic output directed back onto that same sensory neuron (**Figure S9b**). For all pairs, a single sensory neuron contributed only a small fraction of a PSI-IhN’s total sensory input and received only a small fraction of that interneuron’s axoaxonic output (**Figure S9c-g**). This indicates that individual PSI-IhNs integrate convergent input from many sensory neurons of the same class and distribute their axoaxonic synapses broadly across many afferent terminals, mainly of the same class. Consistent with this organization, the axons of PSI-IhNs often extend much farther along the rostral-caudal axis than their dendrites (**Figure 4c**). These features support a circuit architecture in which subtype-selective PSI is implemented predominantly through lateral inhibition. This is consistent with previous findings that disrupting GABAergic axoaxonic synaptic transmission resulted in large, expanded receptive fields of spinal cord neurons that respond to light touch of the skin (Chirila et al., 2022).

Previous studies defining spinal cord IhN populations based on neurochemical identity have also identified molecularly distinct interneurons that mediate PSI (Boyle et al., 2019; Cui et al., 2016; Davis et al., 2023). We therefore asked whether the connectivity-defined PSI-IhN populations correspond to one or more molecularly defined classes. We focused on PV^+^ GABAergic interneurons, which have been proposed to mediate PSI of myelinated LTMRs (Boyle et al., 2019; Hughes et al., 2012). To examine this, we generated an EM dataset from *PV^Cre^; Rosa26^LSL-Matrix-dAPEX2^* mice, in which PV^+^ interneurons were labeled with an EM reporter (**Figure S10a-c**). Consistent with prior reports, PV^+^ IhN axons frequently formed presynaptic contacts onto sensory neurons, although some PV^+^ axons did not form axoaxonic synapses within the imaged volume. Thus, at least some PV^+^ interneurons are indeed PSI-IhNs.

To identify the sensory neuron subtypes connected to PV^+^ PSI-IhNs, we developed a classifier that operates directly on 2D EM images. This approach enables sensory neuron subtype identification from a single EM section without requiring full cell segmentation, albeit with slightly reduced accuracy compared to the SPINE classifier (**Figure S10d**). Using this classifier, we analyzed 44 sensory neurons whose axons are postsynaptic to APEX-labeled PV^+^ axons. Nearly all postsynaptic sensory neurons identified were predicted to be myelinated LTMRs, with 28 classified as Aδ-LTMRs and 11 as Aβ-LTMRs (**Figure S10e**). We also examined sensory inputs onto PV^+^ neuron dendrites and found that most inputs were likewise predicted to arise from A-LTMRs, with 34 of 53 classified as Aδ-LTMRs and 10 classified as Aβ-LTMRs (**Figure S10f**). These results indicate that most PV^+^ interneurons both receive input from and preferentially inhibit Aδ-LTMRs, suggesting their identity as Islet-PSI-IhN^Aδ-LTMR^. Notably, both PV^+^ PSI-IhNs and Islet-PSI-IhN^Aδ-LTMR^ neurons have myelinated axons (**Figure S10g,h**), supporting the idea that these PV^+^ IhNs are a specialized subset of PSI-IhNs that mediate fast PSI of fast conducting LTMRs.

Taken together, our findings suggest that sensory neuron evoked GABAergic PSI operates largely within parallel, subtype-specific homotypic pathways. Distinct sensory neuron classes recruit connectivity- and molecularly-defined PSI-IhN populations, enabling selective homotypic regulation of sensory gain and shaping receptive field properties of spinal cord neurons.

### Heterotypic feedforward inhibition (FFI) circuits

Our reconstructions of PSI circuit motifs highlight their largely homotypic organization. However, PSI-IhNs also mediate FFI by forming axodendritic synapses (Boyle et al., 2019) and, in some cases, dendrodendritic synapses (Davis et al., 2023) onto other spinal cord neurons (**Figure 5a**). We therefore sought to examine the wiring logic of the FFI components of the five PSI-IhN populations. Here, we focused on the same reconstructed PSI-IhN populations analyzed above, although we note that additional inhibitory interneuron populations that do not form axoaxonic synapses also contribute to feedforward inhibition and remain to be explored. Importantly, although the reconstructed PSI-IhN neurons form axoaxonic synapses onto sensory neuron axons, each one also forms synapses onto dendrites of spinal cord neurons, and, in fact, most PSI-IhN neuron synapses were found to be axodendritic or dendrodendritic (**Figure 5b**). Therefore, we investigated the sensory neuron synaptic input properties of the spinal cord neurons targeted by each PSI-IhN’s axodendritic and dendrodendritic synapses.

**Figure 5.**
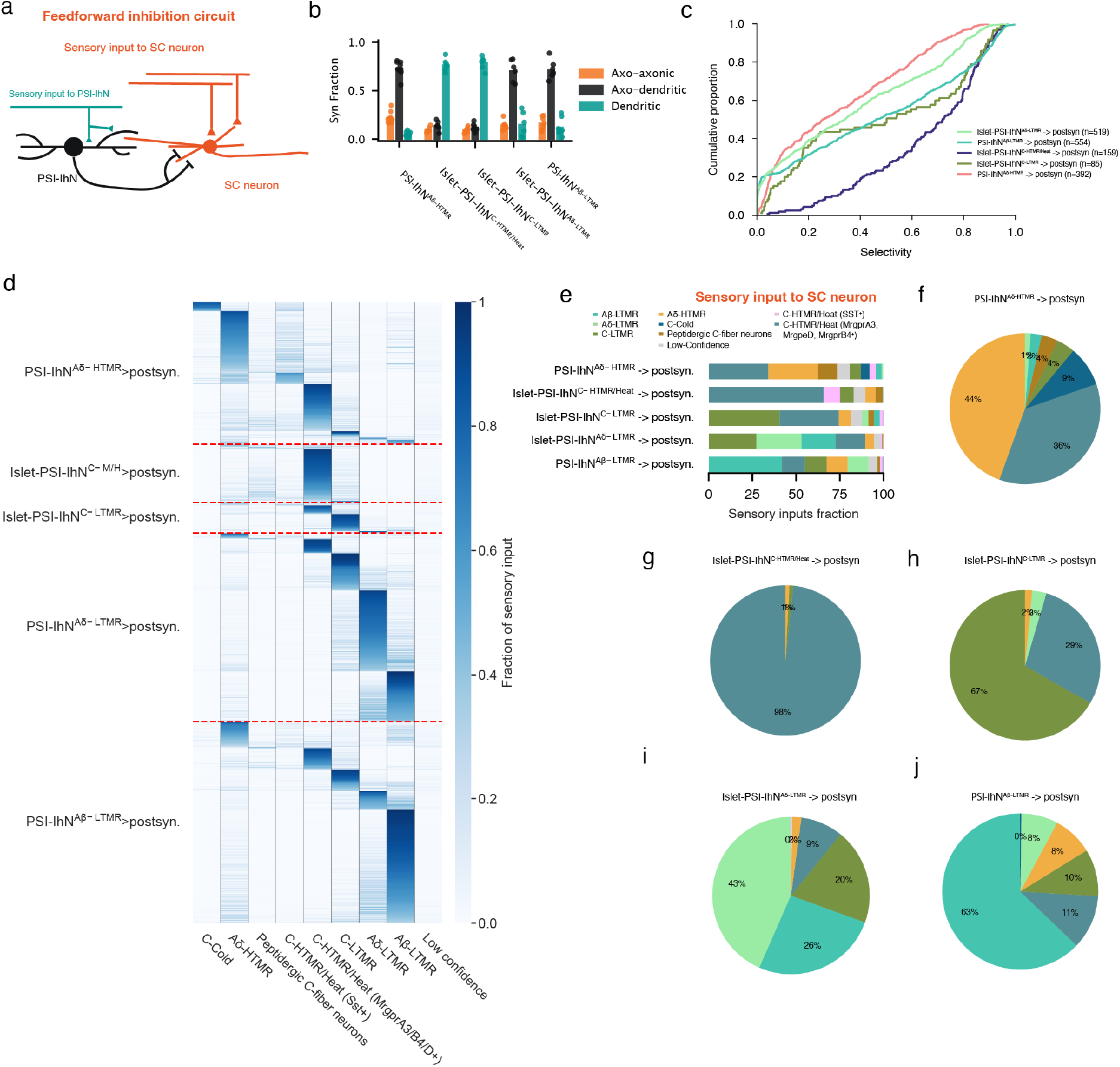
Spinal circuit motifs mediating sensory-evoked heterotypic postsynaptic inhibition. **a**, Schematic of sensory-evoked feedforward inhibition mediated by axodendritic synapses. Sensory neurons of subtype A (light blue) excite GABAergic spinal neurons (red), which in turn inhibit another spinal neuron (dark blue) via axonal output. Because the postsynaptic neuron primarily samples inputs from other sensory subtypes, this motif represents heterotypic feedforward inhibition. **b**, Fractions of axoaxonic, axodendritic, and dendritic output synapses across five PSI-IhN populations. **c**, Cumulative distributions of the homotypic selectivity index for all postsynaptic neurons targeted by axodendritic synapses. Most postsynaptic neurons show low homotypic selectivity, except those postsynaptic to Islet-PSI-IhN^C-HTMR/Heat^ neurons. Numbers of PSI-IhN–postsynaptic neuron connections are indicated. **d**, Fractions of inputs from each sensory subtype onto spinal cord neurons that are postsynaptic to the different PSI-IhNs (rows). (n=392 for PSI-IhN^Aδ-HTMR^ postsynaptic neurons, n=159 for Islet-PSI-IhN^C-HTMR/Heat^ postsynaptic neurons, n=85 for Islet-PSI-IhN^C-LTMR^ postsynaptic neurons, n=519 for Islet-PSI-IhN^Aδ-LTMR^ postsynaptic neurons, n=554 for PSI-IhN^Aβ-LTMR^ postsynaptic neurons.) **e**, Sensory input fractions of postsynaptic neuron populations targeted by axodendritic synapses from each PSI-IhN population. As populations, these postsynaptic neurons sample sensory inputs distinct from those sampled by the presynaptic PSI-IhN populations. **f–j**, Fractions of sensory modules represented by postsynaptic neurons of each PSI-IhN population targeted by axodendritic synapses. (n=254 for PSI-IhN^Aδ-HTMR^ postsynaptic sensory modules, n=121 for Islet-PSI-IhN^C-HTMR/Heat^ postsynaptic sensory modules, n=66 for Islet-PSI-IhN^C-LTMR^ postsynaptic sensory modules, n=426 for Islet-PSI-IhN^Aδ-LTMR^ postsynaptic sensory modules, n=420 for PSI-IhN^Aβ-LTMR^ postsynaptic sensory modules.)

For this, we sampled 1,553 postsynaptic spinal cord neurons contacted by the axodendritic and dendrodendritic synapses of all 34 reconstructed PSI-IhN neurons and used SPINE to infer their direct sensory neuron input composition. Unlike the PSI motifs, which are largely homotypic in nature, the postsynaptic spinal cord neurons receiving FFI from PSI-IhNs often received sensory inputs that differed from those driving the corresponding PSI-IhN neurons (**Figure 5c,d; Figure S11a-e**). For example, while Islet-PSI-IhN^C-LTMR^ neurons as a population receive >90% of their sensory input from C-LTMRs (**Figure 4f**), their axodendritic postsynaptic partner neurons received substantial C-HTMR/Heat input (33.9%) (**Figure 5e**). Similarly, the axodendritic postsynaptic partner neurons of PSI-IhN^Aδ-HTMR^ neurons, as a population, received more synaptic inputs from C-HTMR/Heat neurons than Aδ-HTMRs (**Figure 5e**). The major exception to this general rule was Islet-PSI-IhN^C-HTMR/Heat^ neurons, whose postsynaptic partners largely received a strong enrichment of C-HTMR/Heat input (**Figure 5d,e**). Thus, at the population level, FFI mediated by PSI-IhN neurons is more heterotypic in nature than PSI, and FFI is often skewed toward neurons with abundant C-HTMR/Heat input.

We next asked whether PSI-IhNs mediate FFI onto individual dorsal horn neurons involved in heterotypic sensory pathways. Interestingly, 77.1% of the 1,553 dorsal horn neurons that are postsynaptic to PSI-IhNs had one sensory neuron subtype contributing >50% of their total sensory neuron synapses (**Figure S11f–j; Figure S12g,h**). We therefore used the dominant sensory input to infer the sensory pathway in which each postsynaptic neuron participates. We refer to these spinal cord neurons as sensory modules, with each neuron assigned to the sensory module corresponding to the sensory neuron subtype that contributes >50% of its total sensory neuron synapses (a subset of these modules is characterized in parallel work aimed at defining a superficial dorsal horn circuit wiring diagram (Xiang et al., 2026b)). This framework allowed us to ask which sensory modules are preferentially targeted by PSI-IhN-mediated FFI. Consistent with the population-level analysis of the five PSI-IhN subtypes’ postsynaptic targets, a prominent fraction of the postsynaptic partners of PSI-IhN classes belonged to heterotypic sensory modules. For example, 36% of PSI-IhN^Aδ-HTMR^ postsynaptic sensory modules belonged to neurons within the C-HTMR/Heat modules, and 29% of Islet-PSI-IhN^C-LTMR^ postsynaptic sensory modules also belonged to C-HTMR/Heat module neurons. The sole exception to this general rule was the Islet-PSI-IhN^C-HTMR/Heat^ neurons, which formed synapses almost exclusively onto neurons of the C-HTMR/Heat modules (i.e. homotypic) (**Figure 5f–j**). Notably, for three of the four PSI-IhNs that mediate heterotypic FFI, the most frequently targeted heterotypic module was the C-HTMR/Heat module. Indeed, the PSI-IhN neuronal populations activated by LTMR subtypes and Aδ-HTMRs exhibited biased polysynaptic FFI of the spinal cord neurons that are activated by C-HTMR/Heat sensory neurons. Together, these data indicate that for four of the five GABAergic PSI-IhN populations, feedforward inhibition is substantially heterotypic, with a prominent bias toward FFI of C-HTMR/Heat module neurons.

A third mode of sensory neuron-evoked inhibition engaged by a subset of sensory neuron subtypes is FFI mediated by dendrodendritic synapses (**Figure S12a**). Interestingly, the Islet-PSI-IhN^C-LTMR^ and Islet-PSI-IhN^C-HTMR/Heat^ neurons were found to be unique among the five PSI-IhN populations in forming dendrodendritic synapses (Davis et al., 2023). Individual neurons of these two PSI-IhN populations form thousands of dendrodendritic synapses (**Figure 5b; Figure S12i**), which exceeds the number of their axonal (both axoaxonic and axodendritic) output synapses. We found that the dendrodendritic synapses of these two Islet-PSI-IhN populations form similar FFI motifs as their axodendritic synapse counterparts (**Figure S12c-f**). For Islet-PSI-IhN^C-LTMR^ neurons, 82% of their dendrodendritic targets belonged to C-LTMR module neurons and 17% belonged to C-HTMR/Heat module neurons. In contrast, Islet-PSI-IhN^C-HTMR/Heat^ neurons dendrodendritically inhibited C-HTMR/Heat modules almost exclusively (**Figure S12d-f**). We also noticed that a subset of these dendrodendritic synapses participated in triadic motifs involving a sensory afferent and a feedforward-inhibited spinal neuron (**Figure S12j-m**). To determine how frequently these triads occurred, we quantified, across dSC_APEX datasets, the fraction of sensory axodendritic synapses associated with dendrodendritic inhibition within each glomerulus. Approximately 15% of axodendritic synapses made by C-LTMR and MrgprD⁺ C-HTMR/Heat afferents, and 10% of those made by MrgprA3⁺ C-HTMR/Heat afferents, participated in these triads. Other sensory neuron subtypes rarely formed such motifs, consistent with dendrodendritic synapses being a distinctive feature of Islet-PSI-IhN^C-LTMR^ and Islet-PSI-IhN^C-HTMR/Heat^ neurons (**Figure S12j**).

Thus, while the five PSI-IhN populations predominantly mediate homotypic GABA-mediated PSI through their axoaxonic synapses, four of these IhN populations engage substantial heterotypic FFI to spinal cord neurons. Notably, heterotypic FFI from these PSI-IhNs predominantly targeted C-HTMR/Heat modules. On the other hand, dendrodendritic FFI is limited to two sensory neuron subtypes and is either homotypic or mixed, with the heterotypic form again predominantly targeting C-HTMR/Heat modules. Collectively, PSI-IhNs implement three distinct modes of sensory-evoked inhibition through axoaxonic, axodendritic, and dendrodendritic synapses. These inhibitory circuit motifs provide synaptic substrates through which sensory neuron subtypes regulate sensory gain, spatial acuity, and heterotypic suppression across sensory modalities.

## Discussion

The spinal cord is the first network layer for processing somatosensory signals arriving from the body. More than a dozen somatosensory neuron subtypes convey ensembles of impulses encoding mechanical, thermal, and chemical stimuli to the spinal cord dorsal horn. These signals are integrated within the dorsal horn to generate spinal reflexes and output signals to the brain (Abraira and Ginty, 2013; LaMotte et al., 2014; Woolf, 2011). Yet, compared to initial processing regions of other sensory systems, such as the retina in the visual system, an understanding of the wiring logic of the dorsal horn has been lacking. To resolve this, we generated a large-scale EM volume of the mammalian spinal cord dorsal horn, the dSC1 dataset, together with SPINE, a machine-learning classifier to identify the major somatosensory neuron subtypes, including LTMRs, HTMRs, thermoreceptors, and nociceptors, within this EM volume. These resources enable a systematic analysis of the synaptic organization across major somatosensory neuron subtypes and the shared and subtype-specific circuit motifs that underlie somatosensory information processing within the earliest stage of the somatosensory hierarchy. Our findings reveal a remarkable heterogeneity of somatosensory neuron subtype synaptic architectures and general principles of sensory evoked PSI and FFI for sensory gain control and acuity (**Figure S13**).

Unlike other sensory systems, DRG neurons detect stimuli spanning multiple sensory modalities. Decades of work have described over a dozen primary sensory neuron subtypes, each defined by distinct response properties (Dong and Dong, 2018; Dubin and Patapoutian, 2010; Lallemend and Ernfors, 2012; Qi et al., 2024; Roger et al., 2026), gene expression profiles (Bhuiyan et al., 2024; Sharma et al., 2020; Usoskin et al., 2015; Yu et al., 2024), and morphologies (Abraira and Ginty, 2013; Halata, 1993; Handler and Ginty, 2021; Ikeda et al., 2014; Lumpkin et al., 2010; Qi et al., 2024). These variations are closely linked to the modalities they encode and their specialized functions. By contrast, how central synaptic connectivity of somatosensory neuron subtypes contribute to their specialized functions remains poorly understood. We found that sensory neuron subtypes differ markedly in the organization of their synaptic connectivity. A single Aβ-LTMR forms more than 17,000 axodendritic synapses and distributes information across thousands of spinal cord neurons, far exceeding the number of synapses and postsynaptic partners of HTMR, thermoreceptor, and nociceptor subtypes. This observation suggests that CNS circuits underlying fine, discriminative touch are fundamentally distinct compared to other modalities in how they encode sensory information and broadcast to downstream dorsal horn networks and output neurons. Strikingly, the great majority of LTMR synapses, including their axoaxonic synapses, are located within only a few micrometers of axon terminals, even though individual Aβ-LTMR collaterals and entire Aδ-LTMR and C-LTMR central axons extend over several millimeters. This highly distal arrangement of axodendritic, axoaxonic, and dendrodendritic synapses suggests that individual LTMR branches and terminal arbors may function as semi-independent computational units, capable of recruiting distinct downstream targets.

The five connectomically defined PSI-IhN populations suggest that the dorsal horn uses distinct inhibitory mechanisms to control a sensory neuron subtype’s own gain and to modulate the gain of other sensory subtypes. Homotypic gain control is not distributed evenly across modalities: LTMRs are the primary target of axoaxonic input, in keeping with electrophysiological recordings of robust GABA-mediated PSI of these afferents (**Figure S7**) (Zimmerman et al., 2019), while A-HTMRs and MrgprD⁺ C-HTMR/Heat neurons receive it more sparingly, and thermoreceptors and the Sst⁺/MrgprA3⁺ C-HTMR/Heat populations receive almost none. The sensory neurons of light touch, therefore, are subject to a form of spinal gain control that the sensory neurons encoding heat, cold, noxious stimuli, and itch largely lack. This homotypic gain control is also structurally local: because the PSI-IhN subtypes are preferentially driven by and feed back onto the same sensory class, the resulting circuit motifs implement lateral inhibition within a modality rather than across modalities. FFI, on the other hand, breaks this symmetry. Thus, while PSI stays within a sensory channel, the axodendritic and dendrodendritic outputs of the same PSI-IhNs are prominently heterotypic, reaching postsynaptic neurons driven by other sensory subtypes. The dorsal horn therefore appears to use sensory evoked PSI and FFI for distinct computational roles: PSI tunes an afferent population’s own sensitivity, while FFI mediates suppression of others. Together, these two modes of inhibition offer a synaptic explanation for how gain control, spatial acuity, and cross-modality suppression are implemented by a shared set of interneurons rather than requiring separate circuit motifs for each function.

Interestingly, the PSI-IhNs engage FFI through two types of synapses: axodendritic and dendrodendritic synapses. While axodendritic synapses are abundant across the sensory neuron subtype PSI-IhN modules, dendrodendritic synapses are uniquely associated with the C-LTMR and C-HTMR/Heat modules and often form triadic synaptic arrangements (Davis et al., 2023; Ribeiro-da-Silva et al., 1985). Notably, dendritic outputs constitute the predominant class of output synapses made by Islet-PSI-IhN^C-LTMR^ and Islet-PSI-IhN^C-HTMR/Heat^ neurons. Similar synaptic triads have been described in the visual thalamus, where they are proposed to provide rapid, spatially localized inhibition tightly coupled to sensory drive (Crandall and Cox, 2012; Guillery and Sherman, 2002; Sherman, 2004). Future work defining the physiological properties of the C-LTMR and C-HTMR/Heat triads and the receptors they use will help to understand the computations implemented by this dorsal horn inhibitory motif.

It is noteworthy that LTMR subtypes, which convey light touch signals, and nociceptor subtypes, which encode noxious stimuli, mainly impose PSI-IhN-mediated feedforward inhibition of C-HTMR/Heat modules, which encode medium forces acting on the skin. Therefore, we speculate that heterotypic FFI mediated by PSI-IhN populations may sharpen force tuning input/output relationships in the dorsal horn, whereby light touch pathways mediated by LTMRs suppress medium force threshold mechanosensory input pathways. Similarly, high force mechanoreceptors that convey noxious forces also suppress the medium force threshold pathways (**Figure S13c-e**). This organization may also help to explain how mechanoreceptor activation suppresses itch-conveying signals from C-HTMR/Heat (MrgprA3^+^, MrgprD^+^, and Sst^+^) neurons and thus itch perception. Finally, while we focus on the remarkable range of somatosensory neuron subtype synaptic architectures and principles of sensory evoked inhibition, the dSC1 dataset and SPINE classifier will help resolve additional dorsal horn circuit motifs and cell types underlying somatosensation and how the dorsal horn may be reorganized in tactile and thermal over-reactivity disorders and states of chronic pain and itch.

**Figure S1.**
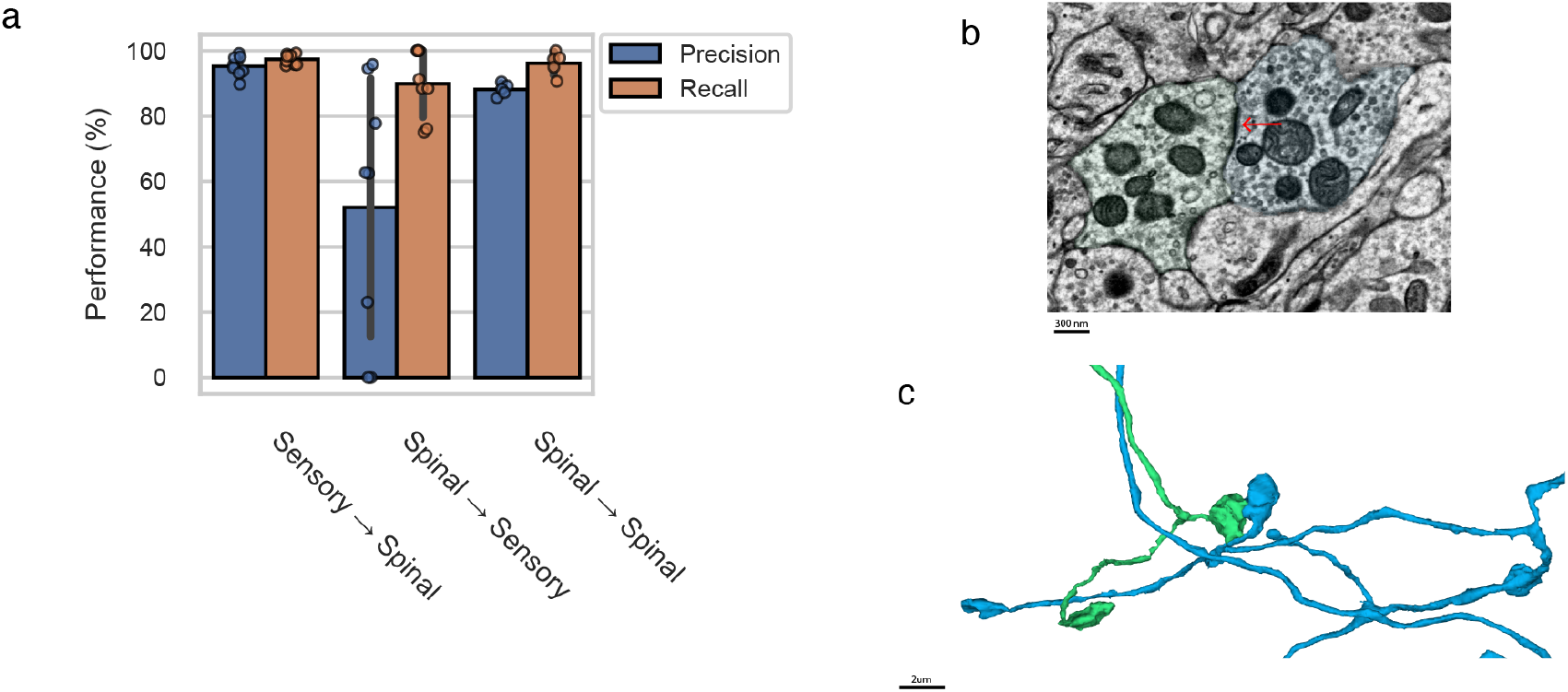
Validation of synapse predictions. **a**, Precision and recall for different synapse-type predictions (**Methods**). The synapse prediction model has a low error rate when predicting axodendritic synapses formed by sensory neurons onto spinal cord neuron dendrites, as well as axodendritic synapses formed by spinal cord neurons onto other spinal cord neurons. However, it shows a higher false positive rate when predicting axoaxonic synapses made onto sensory neurons, especially for C-Cold neurons and peptidergic C-fiber neurons. **b**, Example of a false positive axoaxonic synapse prediction on C-Cold neurons. Two adjacent C-Cold neurons form a symmetric membrane density that may confuse the synapse prediction model. **c**, 3D view of the two adjacent C-Cold neurons.

**Figure S2.**
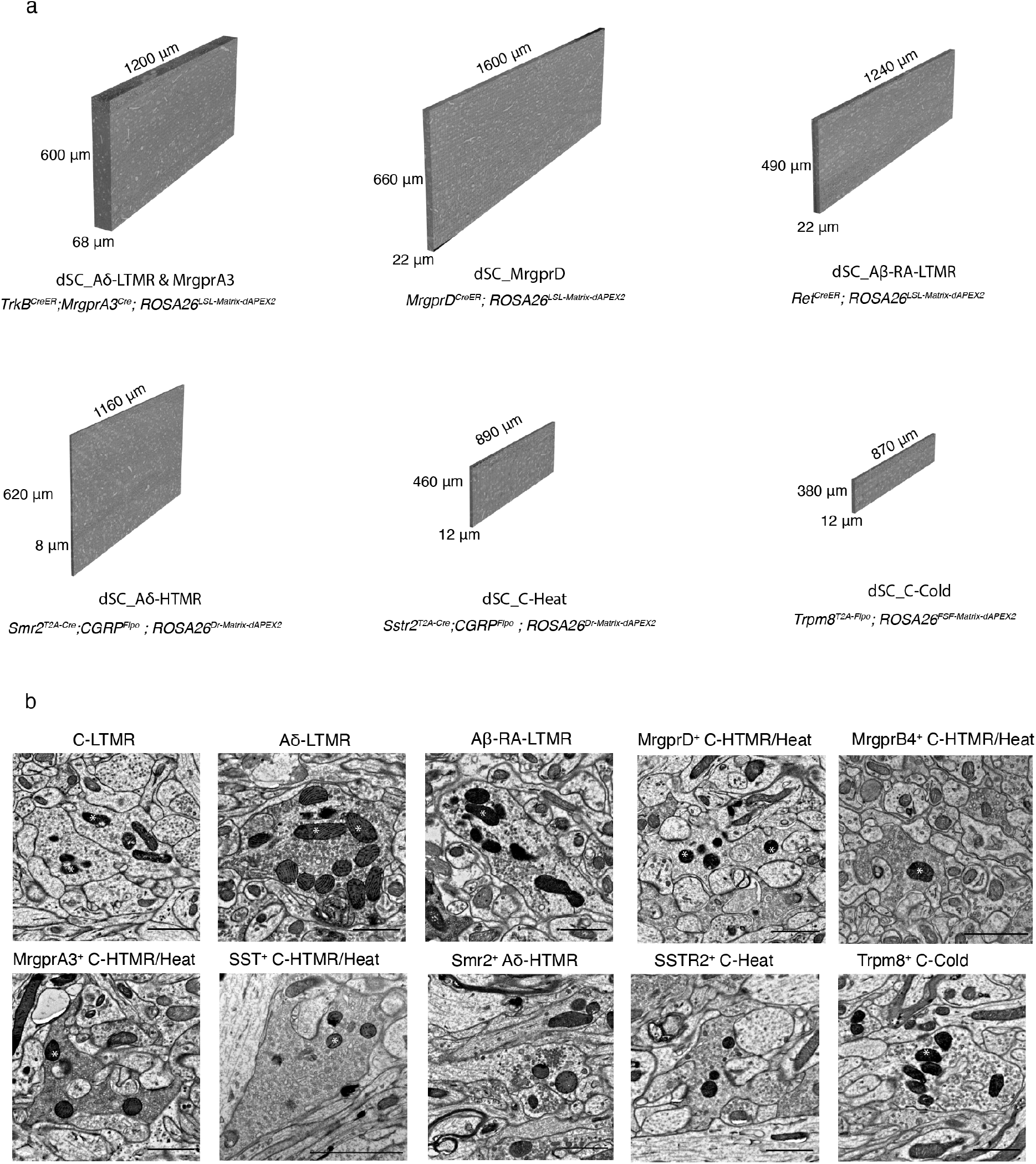
dSC_APEX datasets with genetically labeled sensory neuron subtypes. **a**, Six EM volumes in which different sensory neuron subtypes are genetically labeled, illustrating the diversity of sensory inputs sampled in this study. C-LTMRs, Aβ-LTMRs, and MrgprD⁺ C-HTMR/Heat neurons were labeled by crossing a *ROSA26^LSL-Matrix-dAPEX2^* EM reporter line to *Th^T2A-CreER^*, *Ret^CreER^*, and *MrgprD^CreER^* mice, respectively. Smr2⁺ Aδ-HTMRs (myelinated mechano-nociceptors) and Sstr2⁺ C-Heat neurons (thermo-nociceptors) were labeled by crossing the *ROSA26^DR-Matrix-dAPEX2^*mouse line to *Smr2^T2A-Cre^; CGRP^Flpo^* and *Sstr2^T2A-CreER^; CGRP^Flpo^* mice, respectively. C-Cold neurons were labeled by crossing the *ROSA26^FSF-Matrix-dAPEX2^* reporter line to *Trpm8^T2A-Flpo^*mice. Note that Aδ-LTMRs and MrgprA3⁺ C-HTMR/Heat neurons were simultaneously labeled in *TrkB^CreER^; MrgprA3^Cre^*; *ROSA26^LSL-Matrix-dAPEX2^*mice. All dSC_APEX datasets were segmented except dSC_Aδ-HTMR, owing to imperfect image alignment caused by serial sectioning. **b**, EM images of central terminals from genetically labeled sensory neurons in the lumbar spinal cord. MrgprB4^+^ and Sst^+^ C-HTMR/Heat neurons were labeled in single section TEM experiment using *MrgprB4^Cre^*and *Cysltr2^Cre^* animals harboring the *ROSA26^DR-Matrix-dAPEX2^*allele.

**Figure S3.**
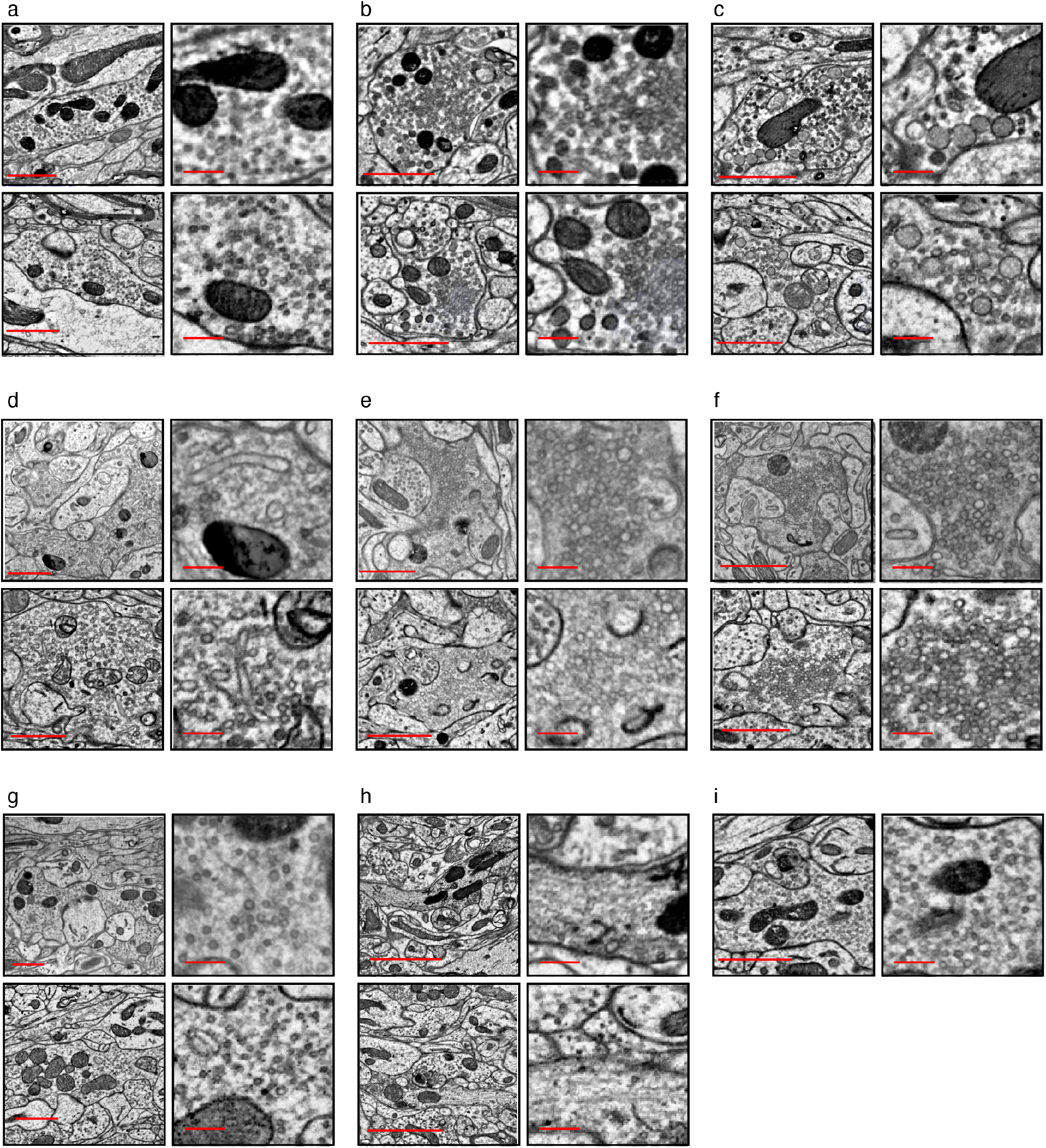
Ultrastructural features of APEX-labeled sensory neurons and the ground truth for training SPINE. **a**, Trpm8⁺ C-Cold neuron morphologies. Top left, APEX labeled Trpm8⁺ C-Cold neuron in SC_C_Cold dataset. Top right, zoomed-in view of the same APEX labeled terminal shown in left. C-Cold terminals contain small, round, clear vesicles that are uniformly distributed within the axonal bouton. Bottom left, C-Cold neuron identified in the dSC1 dataset that was used as ground truth for SPINE training. Bottom right, zoom in view of the same C-Cold neuron shown on the left. **b**, Same as **a** but for peptidergic C-fiber neurons. Top, APEX labeled Sstr2⁺ C-Nociceptor terminals contain dense-core vesicles as well as clusters of small clear vesicles. Bottom, a peptidergic C-fiber neuron identified in the dSC1 dataset similarly has dense core vesicles and is used as ground truth for SPINE training. **c**, Same as **a** but for Aδ-HTMR. Top, Smr2⁺ Aδ-HTMRs labeled by APEX contain large pale vesicles in terminals. Bottom, Aδ-HTMR ground truth annotated in the dSC1 dataset similarly have large pale vesicles. **d**, Same as **a** but for Sst⁺ C-HTMR/Heat neurons. Top, Sst⁺ C-HTMR/Heat neurons labeled by APEX contain characteristic elongated vesicles. Bottom, Sst⁺ C-HTMR/Heat ground truth annotated in the dSC1 dataset share similar ultrastructural features. **e**, Top, MrgprA3⁺ C-HTMR/Heat neurons labeled by APEX and zoomed-in view of the vesicles. Bottom, MrgprD⁺ C-HTMR/Heat neurons labeled by APEX. **f**, Top, MrgprB4⁺ C-HTMR/Heat neurons labeled by APEX and zoomed-in view of the vesicles. Bottom, C-HTMR/Heat (MrgprD⁺, MrgprA3⁺, MrgprB4⁺) ground truth annotated in the dSC1 dataset. C-HTMR/Heat neurons labeled by Mrgpr-family genes share similar ultrastructural features. They have fuzzy cytoplasm and clustered pale vesicles with variable sizes; we thus concatenated all three C-HTMR/Heat neurons as a single cluster. **g**, Same as **a** but for Aδ-LTMR. All three LTMRs contain small round vesicles. **h**, Same as **a** but for Aβ-LTMRs. Aβ-LTMR terminals contain small, round vesicles that are frequently clustered at presynaptic sites, and show abundant neurofilaments (right panels) in the axon terminals. Scale bar, left 2 µm; right, 250 nm. **i**, C-LTMR ground truth examples used to train SPINE. C-LTMRs were APEX labeled in the dSC1 dataset. Scale bar for a-i left panels, 1 µm (except **h**); a-i, right panels, 250nm.

**Figure S4.**
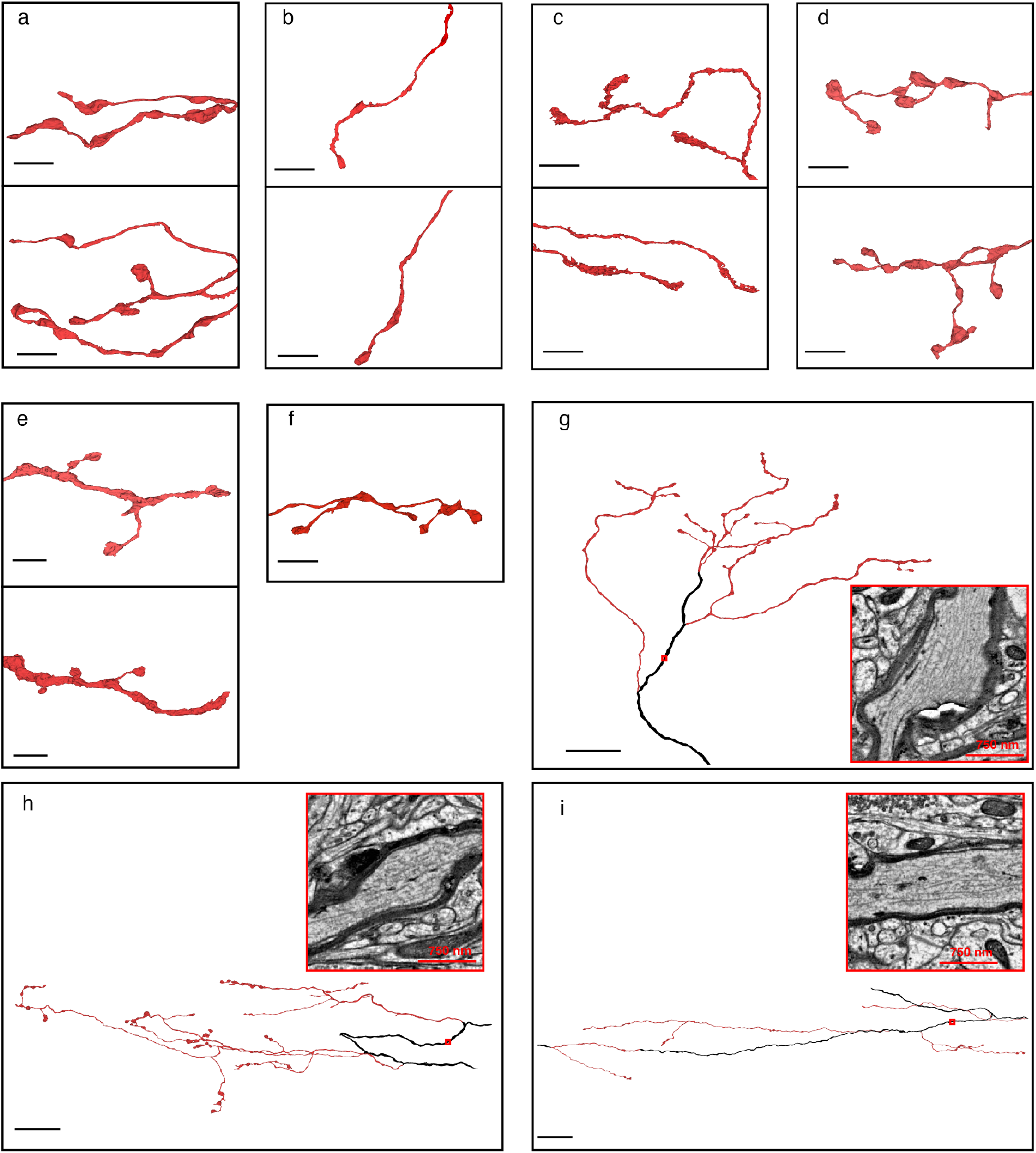
Comparison of 3D morphologies of axonal boutons and cross-dataset validation. **a**, Top, 3D morphology of a Trpm8⁺ C-Cold neuron labeled by APEX. Bottom, C-Cold neuron axonal boutons identified in the dSC1 dataset that were used as ground truth for SPINE training. **b**, Same as **a** but for a peptidergic C-fiber neuron. Top, APEX labeled Sstr2⁺ C-Nociceptor terminals. Bottom, a peptidergic C-fiber neuron identified in the dSC1 dataset used as ground truth for SPINE training. **c**, Top, 3D morphology of a MrgprD⁺ C-HTMR/Heat neuron labeled by APEX. This neuron forms perforated axon terminals. Bottom, 3D morphology of a C-HTMR/Heat (MrgprD⁺, MrgprA3⁺, MrgprB4⁺) ground truth neuron, this neuron similarly has an axon penetrated by postsynaptic components. **d**, Same as **a** but for Aδ-LTMR. Aδ-LTMR axonal boutons are connected by thin neurites and form beads-on-a-string morphologies. **e**, Same as **a** but for Aβ-LTMR. **f**, 3D morphology of a C-LTMR ground truth neuron in the dSC1. Axonal boutons are connected by thin neurites and form beads-on-a-string morphologies. **g**, Myelination pattern for Aβ-LTMRs in the dSC1 dataset. Aβ-LTMRs become unmyelinated (region outlined in red) to form synapses; the myelinated region is colored black. Zoomed-in view show myelin associated with this Aβ-LTMRs ground truth example and abundant neurofilament in the axons. **h**, As in **g,** but for Aδ-LTMR ground truth in the dSC1 dataset. Aδ-LTMRs form beads-on-a-string bouton morphologies and contain fewer neurofilaments. **i**, As in **g**, but for Aδ-HTMR ground truth in the dSC1 dataset. Aδ-HTMRs also contain neurofilaments in their axons. Scale bar **a-f**, 5 µm. **g-i**, 25µm.

**Figure S5.**
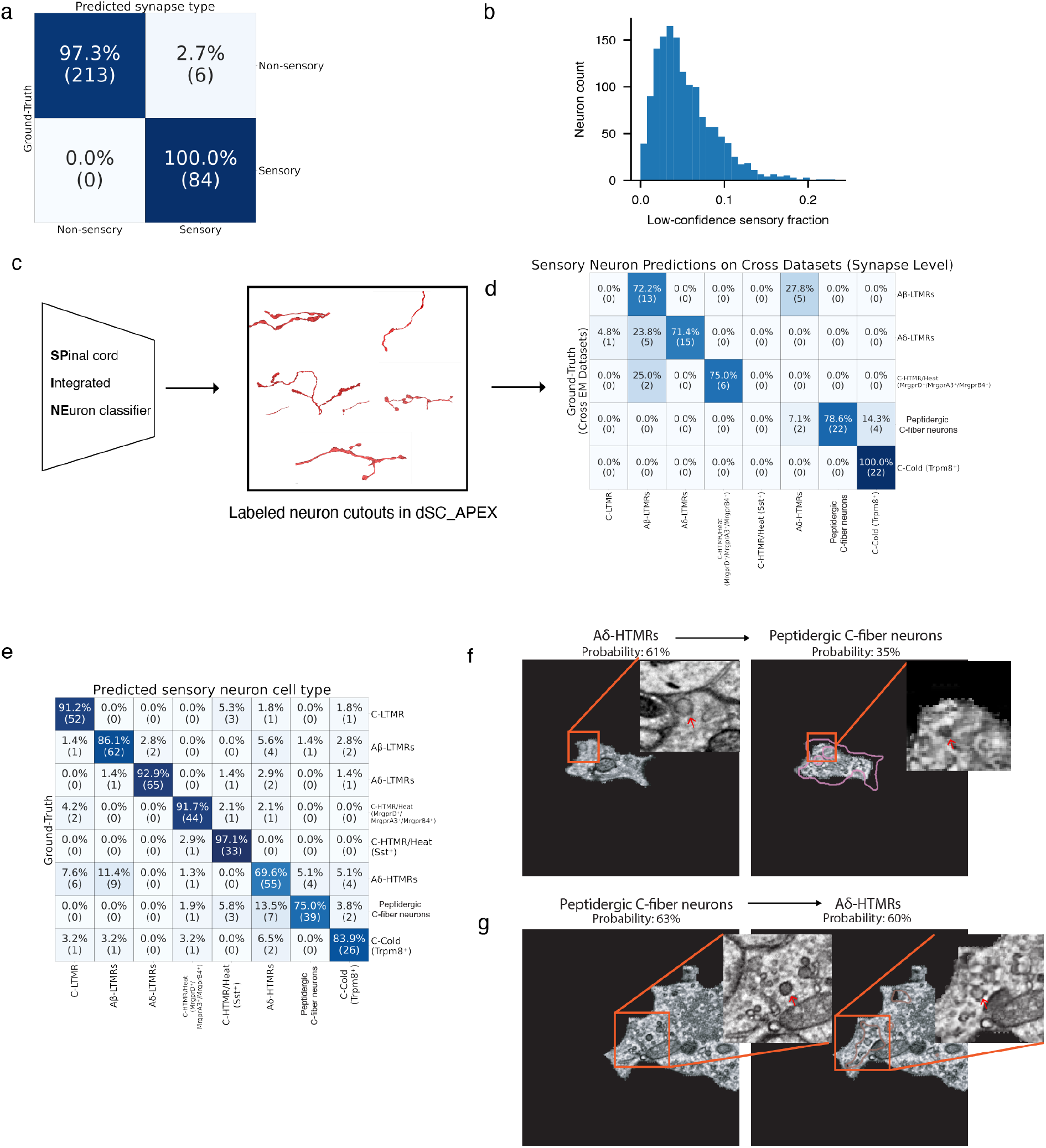
SPINE validation and QuAC feature attributions. **a**, Confusion matrix showing SPINE classification accuracy in stage one: sensory versus non-sensory presynaptic site cutouts. **b**, Distribution of low-confidence sensory predictions across 1,435 neurons classified by SPINE. **c**, Schematic of cross-dataset validation. SPINE was trained on expert-annotated ground truth from the main dSC1 dataset and then evaluated on five additional APEX-labeled, segmented dSC_APEX EM volumes (**Figure S2a**) to assess cell-type generalization across datasets. **d**, Confusion matrix showing SPINE classification accuracy across dSC_APEX datasets. SPINE was not applied to the dSC_Aδ-HTMR dataset because this dataset was not segmented. **e**, Confusion matrix of a 2D classifier trained on dSC1 EM images of sensory neuron subtypes for QuAC. **f**, Example of an Aδ-HTMR image (left) transformed by QuAC into a peptidergic C-fiber-like image (right). Zoomed-in views highlight altered pixels, including conversion of a large pale vesicle into a dark dense-core vesicle. **g**, Reciprocal example showing a peptidergic C-fiber image (left) transformed into an Aδ-HTMR-like image (right), including conversion of a dark dense-core vesicle into a pale vesicle.

**Figure S6.**
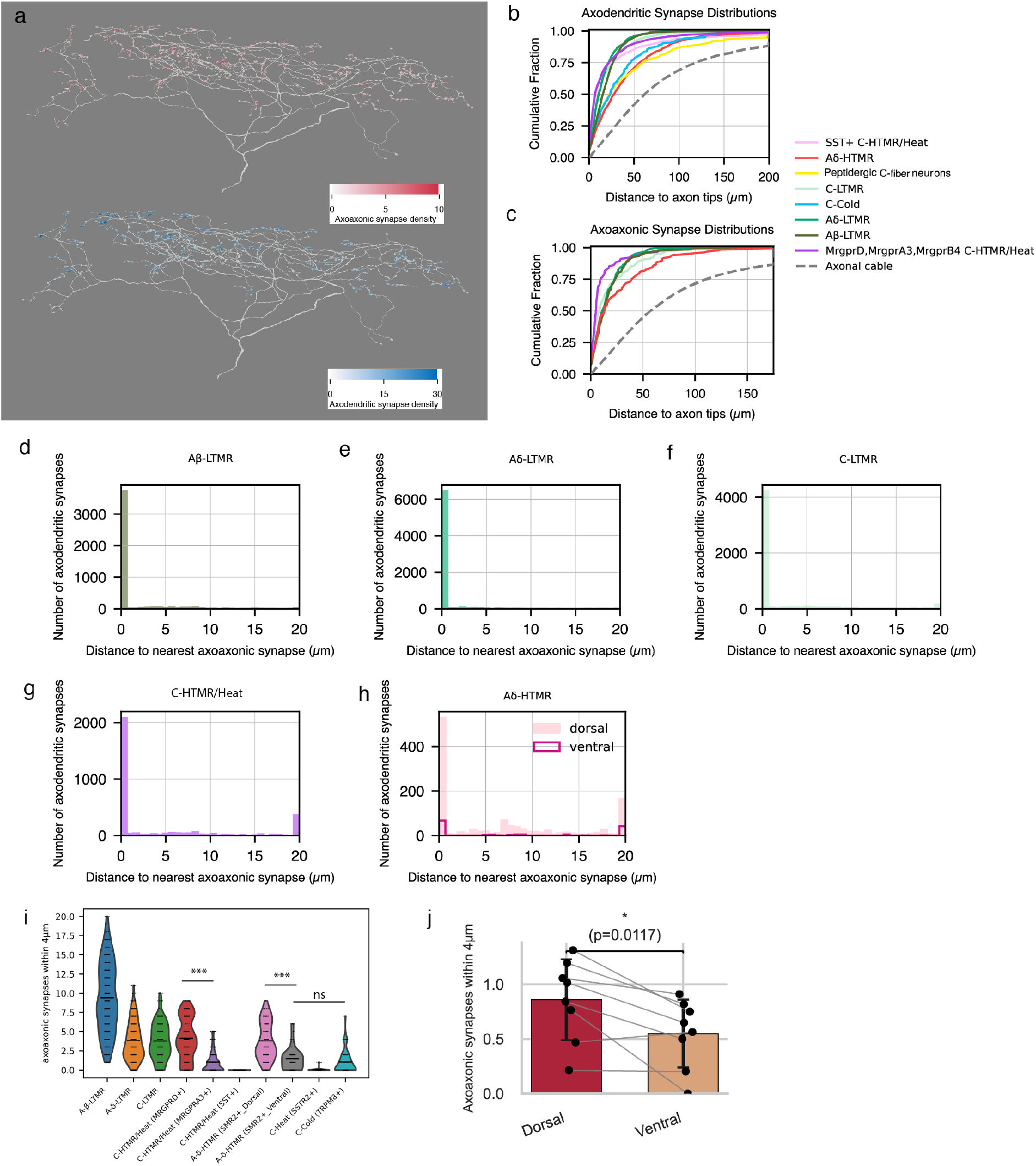
Distribution of axodendritic and axoaxonic synapses. **a**, Example Aδ-LTMR meshes color-coded by axoaxonic synapse density (top) or axodendritic synapse density (bottom). **b**, Cumulative distributions of the cable-length distance from axodendritic synapses to the nearest distal axon tip. Synapses are heavily distributed within the first 25µm from the axonal tips, gray curve shows the percentage of axons within certain distance from axonal tips. n = 10 neurons per subtype, except C-Cold (n = 9). **c**, Cumulative distributions of the cable-length distance from validated axoaxonic synapses to the nearest distal axon tip. n = 2 axons per sensory subtype. **d–h**, Histograms showing the cable-length distance from each axodendritic synapse to its nearest axoaxonic synapse. n = 2 axons per sensory subtype with manually verified axoaxonic synapses. **i**, Number of axoaxonic synapses within 4 μm cable length of each axodendritic synapse in EM volumes in which sensory neurons were genetically labeled. Each dot represents one axodendritic synapse. At least five axons were sampled per genetically labeled sensory neuron subtype. MrgprD^+^ C-HTMR/Heat versus MrgprA3^+^ C-HTMR/Heat, p = 6.38 × 10⁻⁶⁰; Smr2^+^ Aδ-HTMR dorsal versus ventral terminals, p = 5.41 × 10⁻¹²; Smr2^+^ Aδ-HTMR ventral terminals versus Trpm8^+^ C-Cold, p = 1; Kruskal–Wallis test with Dunn’s post hoc test. *n* = 42 Aβ-RA-LTMR, 102 Aδ-LTMR, 45 MrgprD^+^ C-HTMR/Heat, 51 dorsal Aδ-HTMR, 79 C-LTMR, 45 ventral Aδ-HTMR, 42 C-Cold, 118 MrgprA3^+^ C-HTMR/Heat, 34 Sstr2^+^ C-Heat and 44 Sst^+^ C-HTMR/Heat glomeruli. **j**, Complete collateral arbors were reconstructed for 8 Aδ-HTMRs identified in the large EM volume, and the number of axoaxonic synapses within 4 μm cable length of each axodendritic synapse was quantified. Connected dots indicate dorsal and ventral branches of the same Aδ-HTMR. p = 1.17 × 10⁻², Wilcoxon signed-rank test.

**Figure S7.**
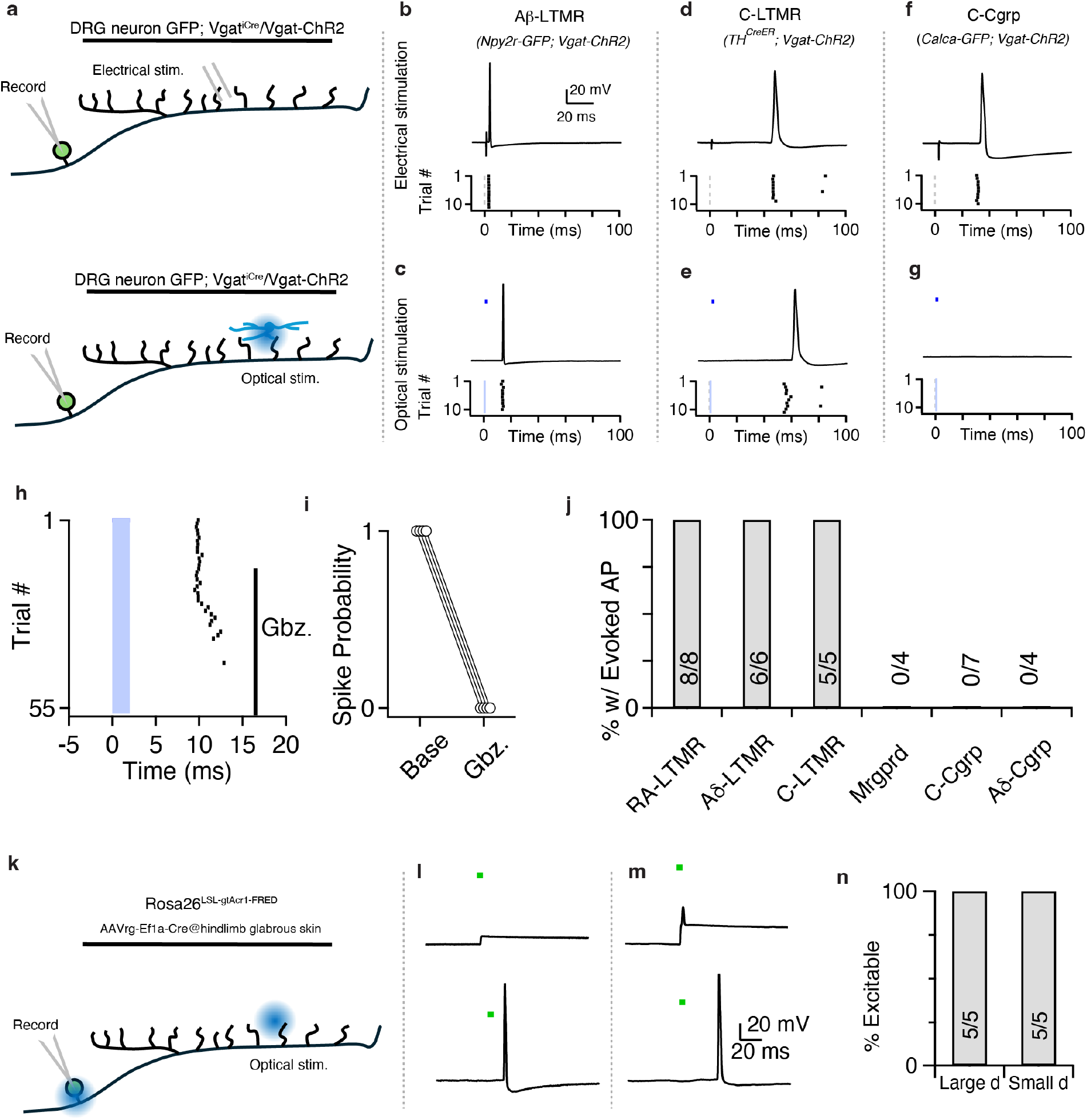
IhN activation generated back-propagated action potentials in LTMRs. **a,** Schematic of experimental setup. The spinal cord and attached DRGs were isolated ex vivo. Whole cell recordings were performed from DRG neuron subtypes identified with reporter mice crossed with *Vgat-ChR2* or *Vgat-IRES-Cre; Rosa26LSL-ChR2* animals to express ChR2 in inhibitory interneurons. Electrical stimuli were delivered to the dorsal horn to confirm connectivity between DRG soma and central terminals, and to identify C, Aδ, and Aβ fibers (top). Optical stimuli were delivered to the dorsal horn through the microscope objective to excite inhibitory interneurons. Experiments were performed in the absence of fast excitatory transmission (AMPARs/NMDARs blocked) (bottom). **b,** Example trace of an identified Aβ-LTMR with action potential back-propagated from the dorsal horn following electrical stimulation (top). Raster of 10 trials of electrical stimulation showing a consistency of the response and low jitter (bottom). **c,** Same neuron as in **b**, but the action potential could be evoked by optical stimulation of dorsal horn inhibitory interneurons. Blue bar indicates light pulse timing. **d-e,** Same as **b-c**, but for a C-LTMR. **f-g**, Same as **b-c**, but for a CGRP-positive C-fiber. Optical activation anywhere in the spinal cord failed to evoke back-propagating action potentials. **h,** Example Aβ-LTMR with action potentials evoked by activation of dorsal horn inhibitory interneurons. Addition of SR95531 (gabazine, Gbz.) to the bath to inhibit GABAARs eliminated evoked action potentials. **i,** Probability of evoking action potentials from optical stimulation of the spinal cord at baseline and after the addition of SR95531 to block GABAARs in four Aβ-LTMRs. **j,** Fraction of neurons in which optical activation of dorsal horn inhibitory interneurons could evoke an action potential for identified subtypes of DRG neurons. Numbers indicate number of neurons. **k,** Schematic of Acr1 excitation in DRG neurons. Same preparation as shown in a, but for animals expressing the light-activated chloride channel Acr1. Hindlimb glabrous skin of Rosa26-LSL-gtAcr1-FRED animals was injected with AAVrg-Cre. Optical stimuli were delivered to the dorsal horn or cell body of randomly recorded DRG neurons. **l,** Putative Aβ-LTMR (identified by A-fiber conduction velocity) with optical activation of the cell body (top) and optical activation of dorsal horn terminals (bottom). Green bar indicates light pulse timing. **m**, Same as in l, but for a C-HTMR (identified by C-fiber conduction velocity and C-HTMR spiking pattern). **n,** Fraction of large (A-fiber) and small (C-fiber) neurons that could be excited by optical activation of dorsal horn terminals. Numbers indicate number of neurons.

**Figure S8.**
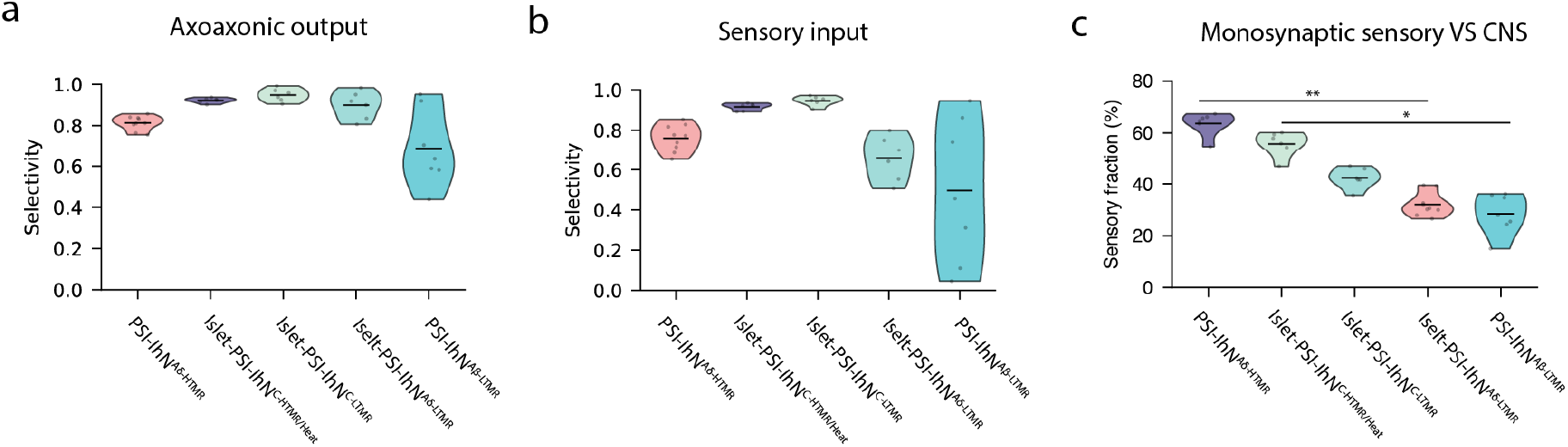
Homotypic feedback inhibition (PSI) mediated by PSI-IhNs. **a**, Fraction of sensory input from the corresponding sensory subtype received by each PSI-IhN population. For example, for Islet-PSI-IhN^C-LTMR^ neurons, selectivity is defined as the fraction of C-LTMR input among all sensory inputs to the neuron. **b**, Fraction of axoaxonic outputs from each PSI-IhN population directed onto the corresponding sensory subtype. **c**, Fraction of direct sensory versus non-sensory (CNS) synaptic inputs onto each PSI-IhN population. (Islet-PSI-IhN^C-HTMR/Heat^ versus PSI-IhN^Aδ-LTMR^, p=2.68×10^-4^, Islet-PSI-IhN^C-LTMR^ versus PSI-IhN^Aβ-LTMR^, p=7.04×10^-3^, Kruskal-Wallis test, Dunn post hoc test).

**Figure S9.**
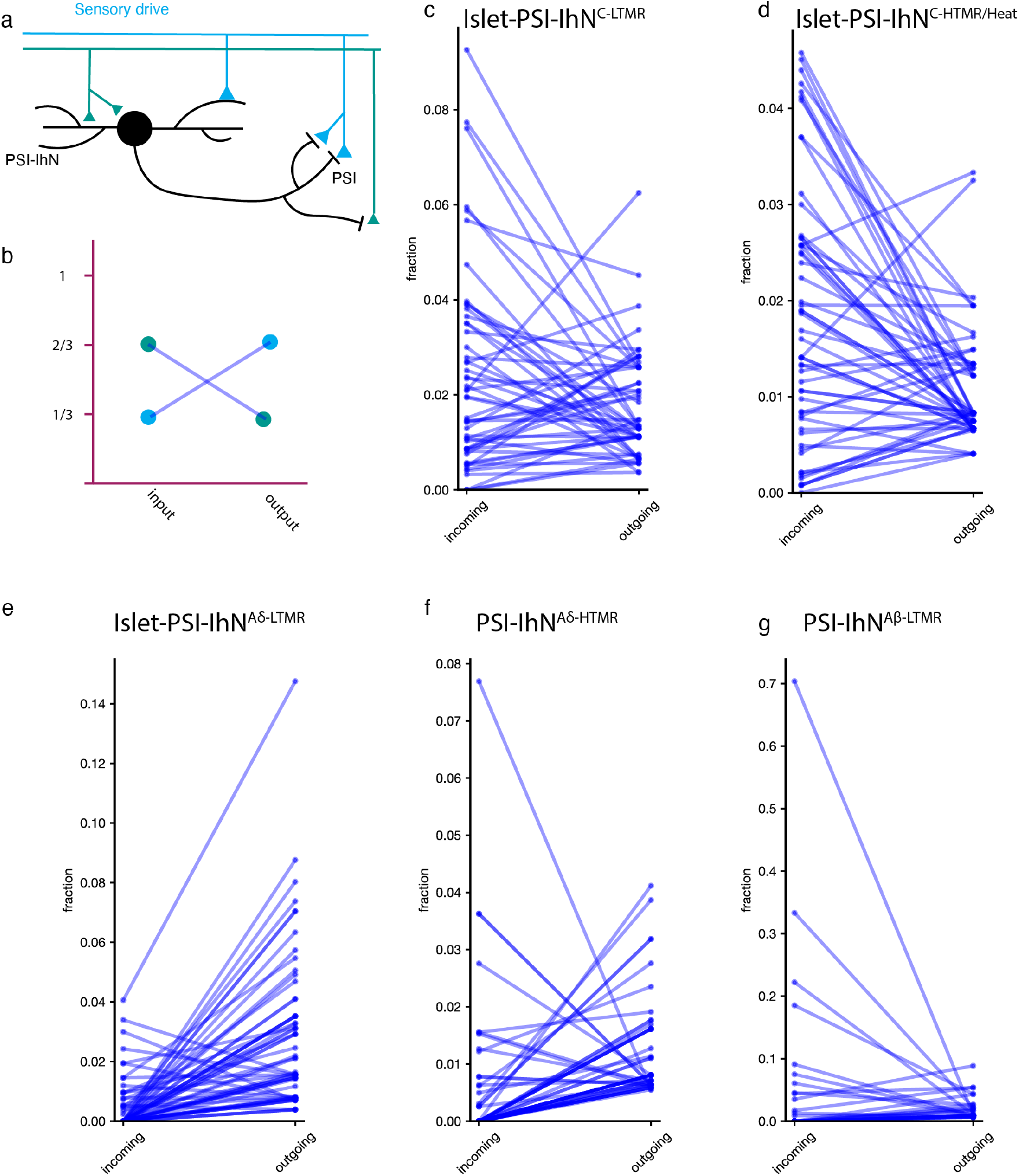
Lateral inhibition mediated by PSI-IhNs. **a**, Schematic of sensory neurons that both drive and receive PSI. A sensory neuron can excite a GABAergic spinal cord neuron that in turn forms axoaxonic synapses onto the same sensory neuron (self-inhibition) or onto other sensory neurons (lateral inhibition). **b**, Quantification of incoming and outgoing fractions for the two example sensory neurons in **a**. The green sensory neuron contributes 2/3 of the sensory input onto the GABAergic neuron and receives 1/3 of its outgoing axoaxonic synapses. The light blue sensory neuron contributes 1/3 of the sensory input and receives 2/3 of the axoaxonic synapses. **c**, Incoming and outgoing fractions for C-LTMR and islet-PSI-IhN^C-LTMR^ pairs. 52 C-LTMR– islet-PSI-IhN^C-LTMR^ pairs were analyzed. **d**, Incoming and outgoing fractions for C-HTMR/Heat and islet-PSI-IhN^C-HTMR/Heat^ pairs. 55 pairs were analyzed. **e**, Incoming and outgoing fractions for Aδ-LTMR and Islet-PSI-IhN^Aδ-LTMR^ pairs. 60 pairs were analyzed. **f**, Incoming and outgoing fractions for Aδ-HTMR and PSI-IhN^Aδ-HTMR^ pairs. 49 pairs were analyzed. **g**, Incoming and outgoing fractions for Aβ-LTMR and PSI-IhN^Aβ-LTMR^ pairs. 31 pairs were analyzed.

**Figure S10.**
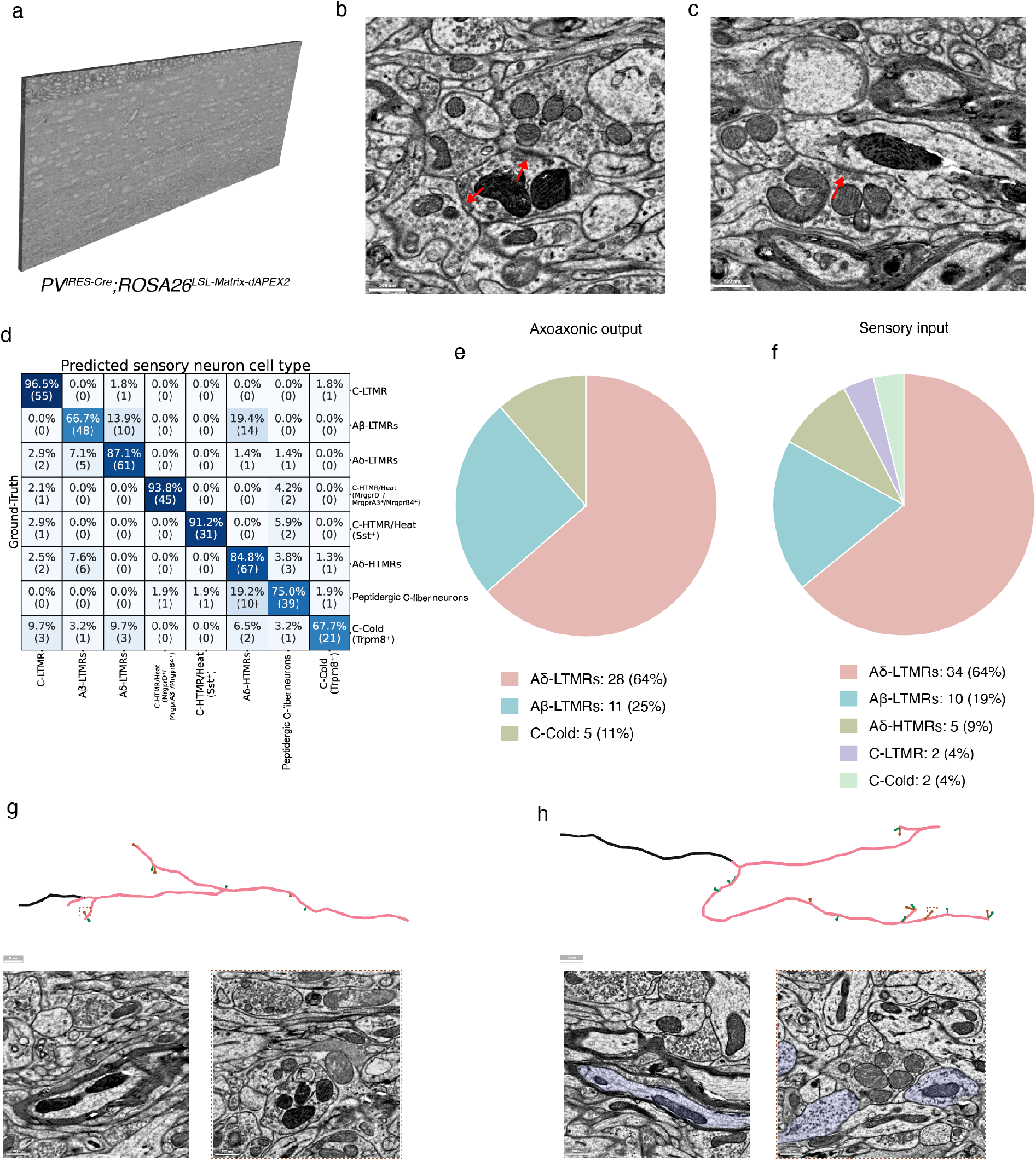
PV^+^ PSI-IhNs form axoaxonic synapses onto A-fiber LTMRs. **a**, An EM volume in which PV^+^ neurons were labeled. The dataset consists of 210 serial-section EM images from a *PV^IRES-Cre^; ROSA26^LSL-Matrix-dAPEX2^* mouse. **b**, APEX-labeled PV^+^ axon forming two axoaxonic synapses onto sensory neurons (red arrows). **c**, Sensory neuron forming axodendritic synapses onto an APEX-labeled PV^+^ dendrite (red arrow). **d**, Confusion matrix of the classifier trained on 4 nm resolution 2D EM images for sensory neuron subtype prediction. **e**, Predicted subtypes of 44 randomly sampled sensory neurons postsynaptic to PV^+^ axons. Most axoaxonic outputs are classified as A-LTMRs, with the majority identified as Aδ-LTMRs. **f**, Same as **e**, but for 53 randomly sampled sensory neurons presynaptic to PV^+^ dendrites. Most inputs are predicted to arise from A-LTMRs, with Aδ-LTMRs comprising the largest fraction. **g**, PV^+^ PSI-IhNs have myelinated axons. Top, skeleton trace of a PV^+^ PSI-IhN axon fragment from the *PV^IRES-Cre^; ROSA26^LSL-Matrix-dAPEX2^* dataset. Black segments indicate myelinated portions of the axon, and pink segments indicate unmyelinated regions. Brown dots mark axoaxonic synapses, and green dots mark axodendritic synapses. Bottom left, EM image showing myelin wrapping the axon. Bottom right, EM image showing an axoaxonic synapse formed by this neuron. **h**, Same as **g**, but from the C-LTMR-labeled dSC1 dataset, showing a myelinated Islet-PSI-IhN^Aδ-LTMR^ neuron.

**Figure S11.**
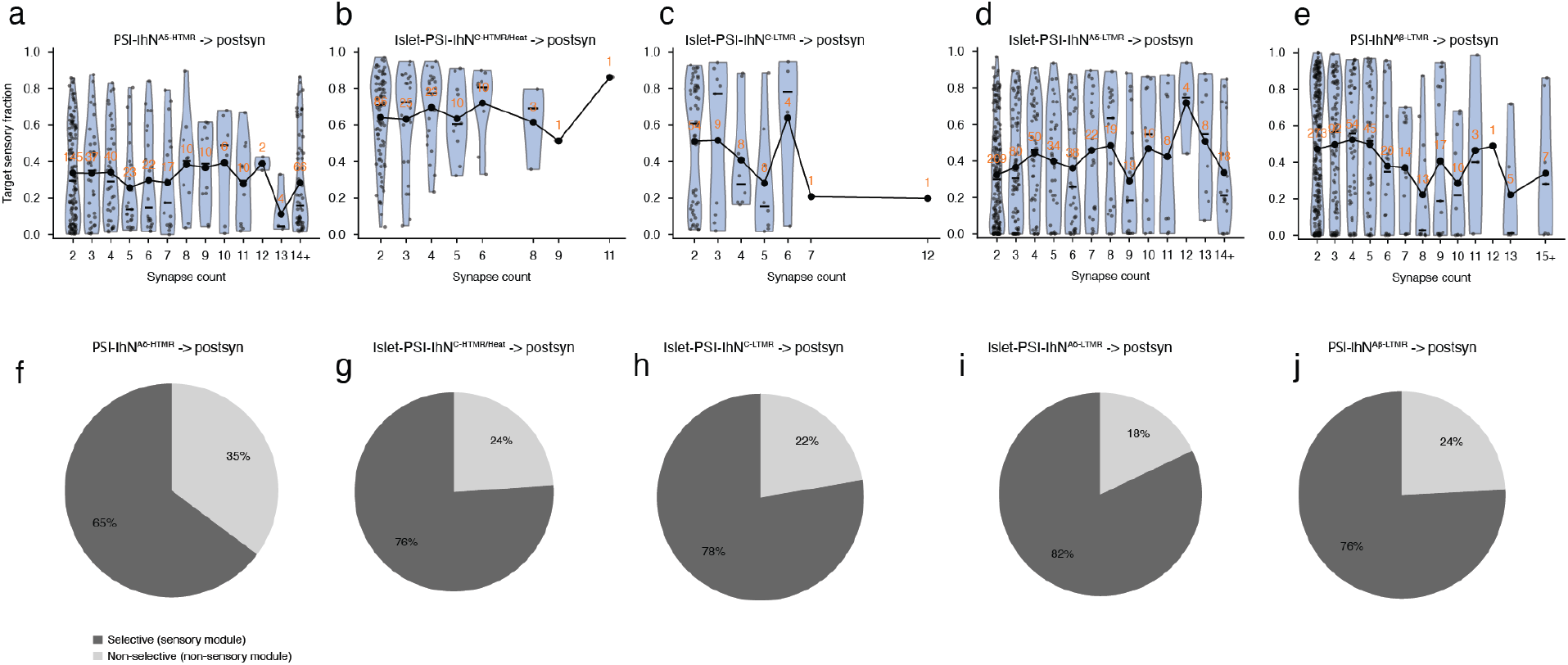
Heterotypic postsynaptic inhibition mediated by PSI-IhNs. **a–e**, Fraction of sensory input from the corresponding sensory subtype onto postsynaptic neurons of each PSI-IhN population targeted by axodendritic synapses. For example, for postsynaptic targets of Islet-PSI-IhN^C-LTMR^ neurons, this value is the fraction of C-LTMR input among all sensory inputs to the postsynaptic neuron. Data are plotted as a function of the number of synapses between the PSI-IhN neuron and the postsynaptic target. Yellow numbers indicate neuron counts. (n=392 for PSI-IhN^Aδ-HTMR^ postsynaptic neurons, n=159 for Islet-PSI-IhN^C-HTMR/Heat^ postsynaptic neurons, n=85 for Islet-PSI-IhN^C-LTMR^ postsynaptic neurons, n=519 for Islet-PSI-IhN^Aδ-LTMR^ postsynaptic neurons, n=554 for PSI-IhN^Aβ-LTMR^ postsynaptic neurons.) **f–j**, Fraction of postsynaptic neurons targeted by axodendritic synapses assigned to specific sensory modules, defined as neurons receiving more than 50% of their sensory input from a single sensory subtype.

**Figure S12.**
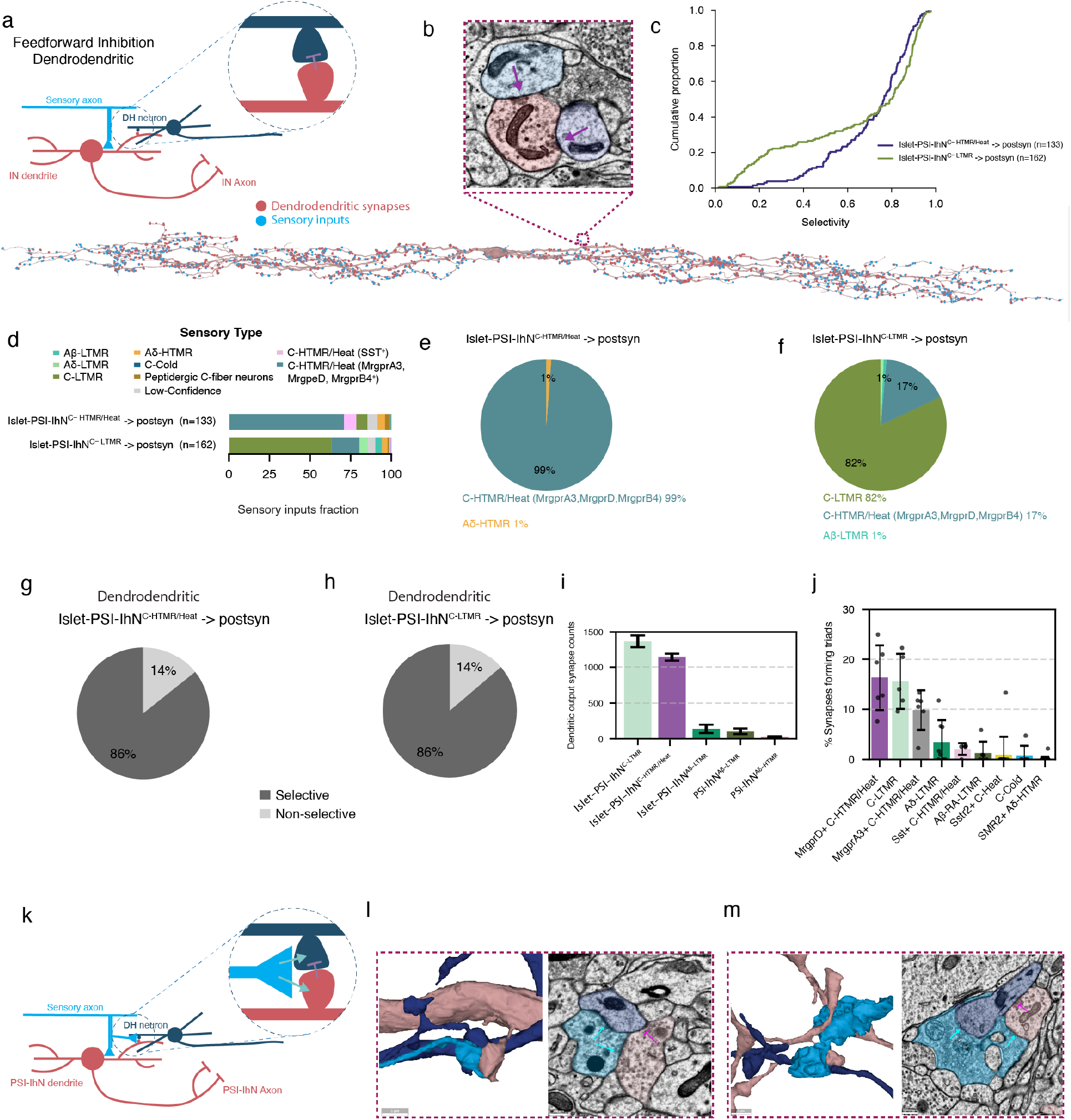
Dendrodendritic synapses mediated by PSI-IhNs. **a**, Schematic of sensory-evoked feedforward inhibition mediated by dendrodendritic synapses. **b**, Example Islet-PSI-IhN^C-HTMR/Heat^ neuron showing sensory input sites and dendrodendritic output synapses. Insets show a dendritic spine forming two dendrodendritic synapses. **c**, Cumulative distributions of the homotypic selectivity index for all postsynaptic neurons targeted by dendrodendritic synapses. Numbers of PSI-IhN–postsynaptic neuron connections are indicated. **d**, Sensory input fractions of postsynaptic neuron populations targeted by dendrodendritic synapses from Islet-PSI-IhN^C-LTMR^ and Islet-PSI-IhN^C-HTMR/Heat^ neurons. **e,f**, Fractions of sensory modules represented by postsynaptic neurons of each PSI-IhN population targeted by dendrodendritic synapses. **g,h**, Fraction of postsynaptic neurons targeted by dendrodendritic synapses assigned to specific sensory modules. **i**, Number of dendritic output synapses formed by PSI-IhN neurons. **j**, Fraction of axodendritic synapses within each glomerulus that are part of triadic synaptic arrangements. **k**, Schematic of a triadic synapse, in which a sensory bouton contacts the dendrites of two spinal neurons, one of which then forms a dendrodendritic synapse onto the other. **l**, Example triad formed between a C-LTMR bouton and an Islet-PSI-IhN^C-LTMR^ neuron. Axodendritic synapses from sensory neurons are annotated as cyan arrows, dendrodendritic synapses from Islet neurons are annotated as purple line. **m**, same as **i**, example triad formed between a C-HTMR/Heat bouton and an Islet-PSI-IhN^C-^ _HTMR/Heat neuron._

**Figure S13.**
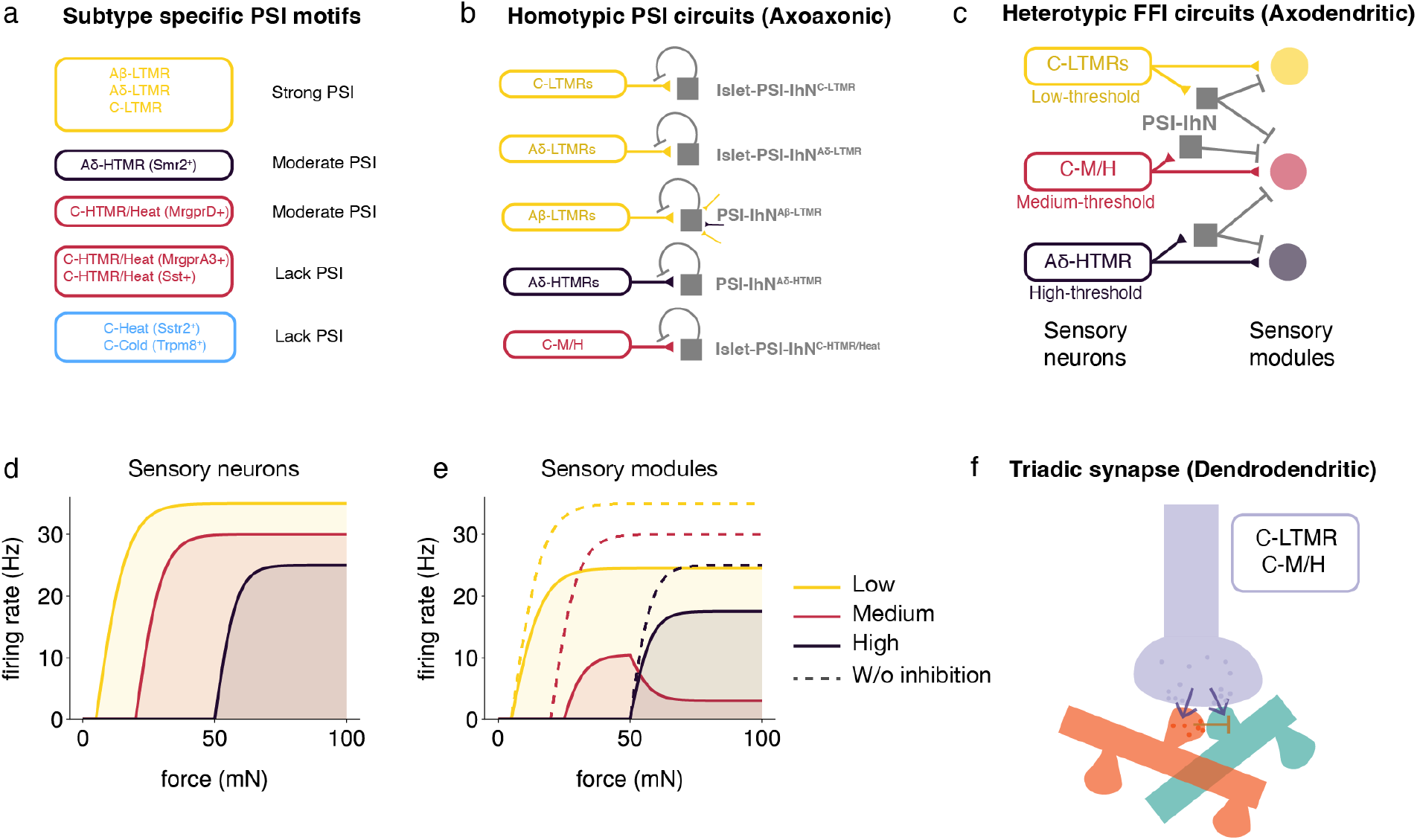
Summary of sensory neuron subtype-evoked feedback and feedforward inhibition motifs of the superficial dorsal horn. **a, b,** Schematic of homotypic feedback inhibition across sensory neuron subtypes. Aβ-LTMRs, Aδ-LTMRs, and C-LTMRs strongly recruit their corresponding PSI-IhN populations and, in turn, receive abundant axoaxonic input from these GABAergic neurons. Aβ-LTMRs also receive a smaller degree of heterotypic PSI recruited by other LTMR subtypes and Aδ-HTMRs. Aδ-HTMR nociceptors and MrgprD^+^ C-HTMR/Heat neurons receive intermediate levels of homotypic feedback inhibition, whereas C-Cold and C-Heat thermoreceptors and Sst^+^ and MrgprA3^+^ C-HTMR/Heat neurons receive relatively sparse axoaxonic input. **c**, Schematic of the heterotypic FFI circuit. C-LTMRs and Aδ-HTMR both inhibit C-HTMR/Heat module through corresponding PSI-IhN populations. This heterotypic inhibition can potentially sharpen modality tuning, as simulated in **d,e**. **d**, Simulated force-tuning curves of C-LTMRs, C-HTMR/Heat neurons, and Aδ-HTMRs, based on data from (Qi et al., 2024); (Lezgiyeva et al., 2026); (Ma et al., 2012); and (Pitcher et al., 2016). **e**, Simulated force-tuning curves of postsynaptic neurons in each sensory module, with and without feedforward inhibition. Without inhibition (dashed curves), postsynaptic tuning largely mirrors presynaptic sensory tuning. With feedforward inhibition (solid curves), the C-HTMR/Heat module shifts to higher response thresholds, loses responses in the LTMR range, and exhibits reduced firing in the nociceptor range. **f**, Schematic of triadic synapses uniquely formed between C-LTMRs or C-HTMR/Heat neurons and their corresponding Islet-PSI-IhN populations.

## Methods

### Mice

All experiments performed in this study were approved by the institutional animal care and use committee of Harvard Medical School. All mice used in the study are of mixed background. For the major C-LTMR labeled dataset, dSC1, a P100 female *Th^T2A-CreER^; ROSA26^LSL-Matrix-dAPEX2^*mouse was used (Abraira et al., 2017; Zhang et al., 2019), tamoxifen (1 mg per day) was administered on P21-P23. For the Aδ-LTMRs and MrgprA3⁺ C-HTMR/Heat labeled dataset, a P50 female *TrkB^CreER^; MrgprA3^Cre^; ROSA26^LSL-Matrix-dAPEX2^* mouse was used, tamoxifen (0.5 mg) was administered on P5 (Han et al., 2013; Rutlin et al., 2014). For the Aβ-RA-LTMR labeled dataset, a P50 female *Ret^CreER^; ROSA26^LSL-Matrix-dAPEX2^* mouse was used, tamoxifen (2mg) was delivered orally to the pregnant mother on E12 (Luo et al., 2009). For the Sstr2⁺ C-Heat neurons labeled dataset, a P66 male *Sstr2^T2A-Cre^; CGRP^Flpo^ ; ROSA26^DR-Matrix-dAPEX2^* mouse was used, tamoxifen (2 mg) was administered on P17 (Qi et al., 2024). For the MrgprD⁺ C-HTMR/Heat neurons labeled dataset, a P60 male *MrgprD^CreER^; ROSA26^LSL-Matrix-dAPEX2^* mouse was used, and tamoxifen (1 mg per day) was administered on P21-P22 (Olson et al., 2017). For the Smr2⁺ Aδ-HTMRs labeled dataset, a P55 female *Smr2^T2A-Cre^; Calca^Flpo^; ROSA26^DR-Matrix-dAPEX2^* mouse was used (Qi et al., 2024). For the C-Cold neuron labeled dataset, a P50 female *Trpm8^T2A-Flpo^; ROSA26^FSF-Matrix-dAPEX2^* mouse was used (Qi et al., 2024). For electrophysiological experiments, described below, mice harboring different combinations of the following alleles were used: *Np2yr-GFP* (Li et al., 2011), *Vgat^IRES-Cre^* (JAX#01962), *TrkB^eGFP^ (Li et al., 2011), TH^CreER^, Vgat-ChR2* (JAX#014548), *Calca-GFP* (GENSAT, RRID: MMRRC_011187-UCD)*, and Mrgprd^eGFP^* (Zylka et al., 2005).

### Tamoxifen administration

Tamoxifen (MilliporeSigma) was dissolved in ethanol (MilliporeSigma) to 10 mg/mL, mixed 1:1 with sunflower seed oil (MilliporeSigma), and vortexed. Ethanol was then removed under vacuum, and the resulting solution was administered to animals by intraperitoneal or oral injection.

### Sample preparation

Mice were transcardially perfused with a fixative (2% paraformaldehyde and 2.5% glutaraldehyde in 4 mM CaCl₂, 0.15 M sodium cacodylate buffer (pH 7.4)). Parasagittal slices (200–300 µm) of the L3–L4 lumbar spinal cord were cut on a vibratome, rinsed in cacodylate buffer, and processed with a 3,3′-diaminobenzidine (DAB) reaction. An enhanced reduced osmium–thiocarbohydrazide–osmium (rOTO) heavy-metal staining protocol was then applied, followed by dehydration through a graded ethanol series and embedding in LX-112 resin (Ladd Research Industries). Samples were polymerized at 60 °C for 3 days. The step-by-step staining protocol has been described previously in (Zhang et al., 2019).

### EM volume acquisition and imaging

Seven spinal cord dorsal horn EM volumes were collected and imaged using the GridTape pipeline for automated serial-section TEM (Phelps et al., 2021). Embedded samples were cut (∼45 nm) with a diamond knife on a modified ATUM-ome and collected onto Luxel GridTape. For the major C-LTMR–labeled dataset, dSC1, 4,105 serial sections were collected with no section loss. After section 3,109, ten sections—3141, 3159, 3171, 3258, 3806, 3858, 3889, 3967, 3968, and 3969—had only half of the region of interest (ROI) landing on the transparent slots. For the other datasets, 170 to 1,499 serial sections were collected without section loss.

Following GridTape sectioning, imaging was performed on an automated TEM system. In total, ∼880 TB of 16-bit raw data were acquired over approximately one year. For the major dataset, nine sections (77, 102, 114, 116, 128, 152, 154, 161, 167) were damaged during imaging. Raw images were converted to 8-bit and stitched into contiguous three-dimensional volumes using an elastic spring-mesh algorithm (AlignTK).

### Segmentation and synapse prediction

The entire C-LTMR–labeled volume, dSC1, was segmented, and automated synapse prediction was performed to facilitate circuit reconstruction. To generate ground truth for cross-dataset validation of the sensory-neuron classifier, subregions of the other EM volumes were also segmented, except for the Smr2⁺ Aδ-HTMR dataset (see “Sensory neuron classifier”).

For segmentation, EM image alignment was first refined using a self-supervised convolutional neural network to apply localized non-rigid warps around imaging artifacts. To reduce merge errors, a separate network was trained to detect myelin; myelin-positive regions were masked prior to neuronal segmentation. We then applied convolutional neural networks to segment the dataset into neuronal fragments (excluding masked myelin regions). For synapse prediction, a 3D convolutional network was trained to classify synaptic-cleft voxels. Cleft probability maps were thresholded and segmented by connected components. A separate network was trained for partner assignment, predicting presynaptic and postsynaptic partner voxels. This model was run once per cleft, and the centroids of the predicted partner masks were recorded as the presynaptic and postsynaptic coordinates associated with each cleft (Lee et al., 2017).

Automated segmentations were ingested into the ChunkedGraph and Connectome Annotation Versioning Engine (CAVE) for proofreading and annotation (Dorkenwald et al., 2023). Synapse predictions were imported (known as synapses_version3 on CAVE) so that corresponding segmentation objects linked to each annotation were dynamically updated during proofreading. This setup enabled queries for an up-to-date connectivity graph and associated metadata, including cell-type annotations.

To assess the accuracy of automated synapse predictions, we manually reviewed reconstructions from eight sensory neurons, one from each of eight subtypes (C-LTMRs, Aδ-HTMRs, Aβ-LTMRs, Aδ-LTMR, peptidergic C-fiber neurons, MrgprA3⁺, MrgprB4⁺, and MrgprD⁺ C-HTMR/heat neurons, Sst^+^ C-HTMR/heat neurons, and C-cold neurons), and five spinal interneurons. For each sensory neuron we inspected every predicted axodendritic (sensory-to-spinal) and axoaxonic synapse (spinal-to-sensory), and for each spinal interneuron we inspected every predicted axodendritic (spinal-to-spinal) synapse, classifying each prediction as a true or false-positive synapse, and separately checking for synapses the algorithm had missed (false negatives). In total, 2,216 predicted synapses were reviewed (1,334 sensory axodendritic, 294 sensory axoaxonic, 588 spinal axodendritic), alongside 1,932 confirmed true synapses (1,311, 196, and 425, respectively). Precision was defined as the fraction of predicted synapses that were true positives (1 − false positives / total predictions checked), and recall was defined as the fraction of true synapses that were correctly predicted (1 − false negatives / (true positives + false negatives), where true positives = total predictions − false positives).

### Generating ground truth for sensory neuron subtypes

To establish ground truth labels, three expert neuroanatomists (W.X., Q.Z., S.C.) annotated sensory axonal boutons in the major C-LTMR labeled EM volume, dSC1, guided by laminar position and ultrastructural features. Criteria were informed by prior light-microscopy mapping of central termination patterns and APEX labeling experiments. Sensory boutons were distinguished from CNS axons by their large terminals forming complex glomerular arrangements. Subtype definitions were as follows:

1. C-Cold (Trpm8⁺) — Terminate in the most dorsal region of Lamina I, just beneath the white matter. Boutons contain uniformly distributed round, clear vesicles with little to no dense-core vesicles (DCVs).
2. Peptidergic C-fiber neurons (CGRP⁺; includes Sstr2⁺ C-Heat neurons) — Terminate in lamina I/IIo. Boutons exhibit numerous dark DCVs and clusters of small round vesicles.
3. Aδ-HTMRs — Terminate in lamina I/IIo with collaterals extending into Laminae III–IV. Boutons contain many dark DCVs plus occasional very large, pale round vesicles; axons have larger diameters compared to peptidergic C-fiber neurons and are myelinated.
4. C-HTMR/Heat (Sst⁺) neurons — Terminate in lamina IIo/IIid. Boutons are characterized by large, elongated vesicles.
5. C-HTMR/Heat (MrgprD⁺/MrgprA3⁺/MrgprB4⁺) neurons — Terminate in Lamina IIid. Because the ultrastructural features of MrgprD⁺, MrgprA3⁺, and MrgprB4⁺ neurons are highly similar and difficult to distinguish, these three neurons were grouped into a single class for classification. Boutons exhibit fuzzy cytosol and dense clusters of pale, round vesicles with variable sizes; their endings correspond to classic type 1 glomeruli.
6. C-LTMRs — Terminate in Lamina IIiv. Boutons contain round, clear vesicles and lack the clustered vesicle pattern typical of type 1 glomeruli. (These neurons are APEX-labeled in dSC1 datasets.)
7. Aδ-LTMRs — Terminate primarily in Lamina IIiv–III. Boutons contain small, round, clear vesicles that are uniformly distributed and resemble type 2 glomeruli. Compared with C-LTMRs, they exhibit more and larger mitochondria. Terminals are typically spherical and linked by thin inter-bouton processes, yielding a “beads-on-a-string” morphology. Their axons are myelinated.
8. Aβ-LTMRs — Terminate in Laminae IIiv–IV. Boutons contain small, round, clear vesicles with abundant neurofilaments that often displace vesicles from the center. Terminals are large and rod-like rather than spherical. Axons are myelinated.

### SPINE - SPinal cord Integrated NEuron classifier

The SPinal cord Integrated NEuron classifier (SPINE) is a two-stage classification pipeline built to predict sensory neuron subtypes from local electron microscopy ultrastructure and morphology at the level of individual synapses. The pipeline consists of two sequential stages: (1) a binary sensory vs. non-sensory classifier, and (2) a downstream 8-class sensory neuron subtype classifier.

For stage one, we extracted a local 3D EM cutout centered directly on the presynaptic site. Each cutout was sampled at a voxel resolution of 32 x 32 x 45 nm, spanning an approximate physical volume of a 4 μm cube. To isolate the ultrastructure of the target axon from neighboring neurons, each cutout was masked using the neuron’s automated segmentation. This masked volume was then embedded using a SegCLR (Dorkenwald et al., 2023) model trained on the MICrONS (Consortium, 2025) dataset to produce a 64-dimensional ultrastructural feature vector. A lightweight, binary Multi-Layer Perceptron (MLP) processed this embedding to classify the synapse as either sensory or non-sensory. Synapses predicted as non-sensory were filtered out, while those classified as sensory progressed to stage 2 of the pipeline.

To predict sensory neuron subtypes, the next stage integrates broader ultrastructural context and local morphology with a 5-10 μm radius surrounding the synaptic bouton. Within the spatial radius, multiple 3D EM cutouts were sampled at 1.5 μm spacing (each maintaining the same 32 x 32 x 45 nm voxel resolution and 4 μm cube volume). As in stage one, each cutout was masked to restrict the input view to only the target axon. Each cutout was embedded independently using two SegCLR models: one trained on the MICrONS dataset and one on the human H01 dataset (Shapson-Coe et al., 2024), resulting in two 64-dimensional embeddings per cutout. For each dataset source, embeddings were aggregated per presynaptic region across all local sampled cutouts by the mean. This generated one MICrONS feature vector and one H01 feature vector. These were concatenated into a unified 128-dimensional ultrastructural representation. Concurrently, the neuron’s 3D mesh geometry was extracted within the same 5-10 μm radius. The local submesh was treated as a graph, where nodes represent mesh vertices in nanometer coordinates. A Graph Neural Network (GNN) (Kipf and Welling, 2017) encoded the spatial topology into a structural embedding vector. The 128-dimension SegCLR vector and GNN structural embedding were concatenated to form the final multi-modal feature vector. This joint representation was fed into a downstream 8-class MLP to predict sensory neuron subtype.

The final class probabilities for the subtype classifier were obtained via a softmax output layer, with predictions assigned to the maximum-probability class. To ensure high-confidence assignments, a confidence threshold was implemented where synapses with a probability below 0.6 were designated as “uncertain”. Both the binary and 8-class MLPs were optimized using the Adam optimizer (Kingma and Ba, 2014) and cross-entropy loss with class-balanced sampling to account for data imbalances. The complete dataset consists of 172 neurons and 2,057 synapses. The data was split at the neuron level where 141 neurons (1,624 synapses) were used for training, and 31 neurons (433 synapses) were held out as an independent test set to evaluate model performance. The corresponding numbers of training and test cutouts, respectively, were 301 and 79 for Aδ-HTMRs, 141 and 31 for C-Cold neurons, 193 and 34 for Sst^+^ C-HTMR/Heat neurons, 207 and 70 for Aδ-LTMRs, 152 and 38 for C-LTMRs, 47 and 19 for unlabeled C-LTMRs, 197 and 48 for MrgprA3/B4/D^+^ C-HTMR/Heat, 231 and 72 for Aβ-LTMRs, and 171 and 52 for peptidergic C-fiber neurons.

### 2D classifiers for Interpretability and Scaled Inference

To facilitate downstream visual interpretability of SPINE and to extend sensory neuron classification to EM datasets lacking automated neuron segmentations, we trained two distinct 2D image-based classifiers. Both models utilized a standard ResNet-18 architecture adapted for 2D grayscale EM input and maintained the exact neuron-level training and testing splits used for the primary SPINE architecture. To investigate ultrastructural features driving classification via Quantitative Attribution with Counterfactuals (QuAC), we trained a 2D classifier restricted to a single slice. This model was trained on 2D EM data sampled at a resolution of 8 x 8 x 45 nm with automated neuron segmentation included in the input. Images were 500 x 500 pixels wide, capturing a context area of 4 μm square centered on the presynaptic region. To classify sensory neuron substypes in new volumes lacking dense neuron segmentations (e.g. the PV^+^ interneuron dataset), we developed a segmentation-free model. This network was trained at a higher native resolution of 4 x 4 x 45 nm using raw EM images with no segmentation mask. To enforce a highly localized focus on the presynaptic active zone, the model utilized a “zoomed-in” 250×250 pixel wide window, providing a tight 1 μm spatial context directly around the presynaptic site. Both models were optimized using the Adam optimizer and cross-entropy loss with class-balanced sampling. Standard 2D data augmentations (random rotations, flips, intensity jittering) were applied during training to improve generalization. Class assignments were generated by taking the argmax of the probability distribution from the final softmax layer.

### Targeted neuron reconstruction and proofreading

Identification of sensory neuron subtypes are described above. All sensory neuron reconstructions reported here were performed in the major C-LTMR–labeled EM volume, dSC1. Within this volume, we targeted sensory axons whose arbors were centered in the EM volume to minimize truncation at borders. Proofreading was carried out primarily in Neuroglancer. Each axon was traced and edited until it exited the EM volume or its entry point at white matter.

We reconstructed: 23 C-Cold neurons (7 partial), 21 peptidergic C-fiber neurons (3 partial), 22 Aδ-HTMRs (4 partial), 18 Sst⁺ C-HTMR/Heat neurons (2 partial), 20 C-HTMR/Heat (MrgprD⁺/MrgprA3⁺/MrgprB4⁺) neurons (4 partial), 16 C-LTMRs (4 partial), 18 Aδ-LTMRs, and 21 Aβ-LTMR collaterals. ‘Partial’ denotes arbors truncated by volume boundaries.

For inhibitory neurons, axons forming axoaxonic synapses onto identified sensory axons were first located and traced retrogradely to their soma and dendrites when possible. Neurons with identified soma were then proofread for downstream analyses. In total, 6 Islet-PSI-IhN^C-HTMR/Heat^, 6 Islet-PSI-IhN^C-LTMR^, 6 Islet-PSI-IhN^Aδ-LTMR^, 9 PSI-IhN^Aδ-HTMR^ and 7 Islet-PSI-IhN^Aβ-LTMR^ were reconstructed.

### Quantification of axoaxonic and axodendritic synapses

To quantify the number of axodendritic synapses associated with entire sensory neurons, we pulled synapses from CAVE in which the sensory neurons were marked as presynaptic and filtered out synapses with clefts smaller than 20 voxels. To quantify the number of axoaxonic synapses, we queried CAVE for synapses in which the sensory neuron was annotated as postsynaptic. We note that our synapse prediction model identified false-positive synapses onto sensory neurons. Therefore, we selected two sensory neurons for each type and manually validated all predicted axoaxonic synapses to remove false-positive predictions.

We also validated our findings by quantifying the number of axoaxonic synapses in other EM volumes in which different sensory neuron subtypes were APEX-labeled. We used the CATMAID collaborative annotation software to manually trace randomly selected axonal boutons (Saalfeld et al., 2009). We reconstructed more than 20 boutons from at least five individual axons for each sensory neuron subtype. We traced the skeletons of the sensory neurons and annotated the axoaxonic synapses formed on the neurons as well as the axodendritic synapses. Synapses were then grouped into individual bouton structures using an iterative 2-µm distance threshold, with merging repeated until no additional synapses could be assigned to a cluster. All synapses within each cluster were considered part of the same glomerulus, and the number of synapses per glomerulus was quantified. The number of glomeruli analyzed for each sensory neuron subtype is indicated in the figure legend.

To quantify the number of spinal cord neurons downstream of each sensory neuron, we queried CAVE for synapses in which the sensory neuron was presynaptic, excluded autapses, retained synapses with clefts of at least 20 voxels as described above, and grouped the remaining synapses by postsynaptic segment. Because the automated segmentation produces many small, unproofread fragments that receive synapses but do not correspond to reconstructed neurons, we counted the level-0 supervoxels agglomerated into each postsynaptic segment and considered an object to be an intact neuron when it comprised at least 40,000 supervoxels. The number of downstream neurons was then defined as the number of distinct intact postsynaptic neurons contacted.

### Synapse-density coloring of neuron meshes

Meshes for selected sensory neurons were retrieved and represented as vertex–face triangulations. Corresponding pre- and postsynaptic centroids were fetched from CAVE and scaled to nanometers. For each neuron, a vertex-adjacency graph was built from the mesh, with edge weights equal to the Euclidean length between neighboring vertices. Each synapse point was snapped to its nearest mesh vertex, and a geodesic (surface) neighborhood of radius R = 4 µm was computed by Dijkstra search on the vertex graph; every vertex within this cutoff received +1 count (multiplied when multiple synapses snapped to the same vertex). Summing across all synapses yielded a per-vertex density map, computed separately for presynaptic and postsynaptic events.

### Skeletonization and synapse–axon tip distance analysis

We generated level-2 skeletons for sensory neurons using the pcg_skel package (Dorkenwald et al., 2025), which leverages the PyChunkedGraph to group all supervoxels of a segmentation into 2 × 2 × 20 μm level-2 chunks (each with a unique ID); the level-2 skeleton was computed over this chunk graph. Synapses were mapped to their containing level-2 chunks and then assigned to the nearest skeleton node.

To quantify distances to axonal tips, skeleton endpoints except the axonal root were taken as axon tips. We computed geodesic (path) distances along the skeleton between each synapse node and all endpoints, recorded the minimum distance per synapse, and plotted the resulting distribution.

### Inhibitory neuron connectivity

Sensory inputs to inhibitory neurons were identified by querying CAVE for synapses onto each inhibitory neuron. For each input, the presynaptic neuron cutout was embedded as described above, and classified as a CNS neuron, a specific sensory subtype, or low-confidence sensory if all sensory subtype scores were ≤ 0.6. For each inhibitory neuron, the proportion of each sensory subtype was computed and normalized to the total number of sensory inputs. To quantify downstream targets of axoaxonic synapses, at least 100 axoaxonic contacts per inhibitory neuron were manually annotated across distinct axon branches. Postsynaptic neuron identities were then determined using the same embedding and classification procedure.

### Lateral inhibition analysis

Connectivity was obtained from CAVE. For each sensory–PSI-IhN subtype pair (e.g. Aδ-HTMR and PSI-IhN^Aδ-HTMR^), we first identified all PSI-IhN cells of the corresponding type and manually split each neuron into axon and dendrite compartments. Using the CAVE synapse tables, we then pulled (1) all synapses from the PSI-IhN axon onto sensory neurons of the chosen subtype, and (2) all synapses from those sensory neurons back onto the PSI-IhN dendrite. For each PSI-IhN– sensory pair, the outgoing fraction was defined as the proportion of all synapses made by that PSI-IhN axon that targeted that specific sensory neuron, and the incoming fraction as the proportion of all synaptic inputs to the PSI-IhN dendrite that originated from that same sensory neuron. If a PSI-IhN axon inhibited a given sensory neuron but we observed no synapses from that sensory neuron back onto the PSI-IhN dendrite, the incoming fraction for that pair was set to zero. Only neuron pairs with an actual PSI-IhN axon → sensory neuron connection were included, and each such connection contributed one incoming and one outgoing fraction value to the analysis.

### Feedforward inhibition with triadic synapses

Inhibitory neuron reconstructions were manually partitioned into dendrites and axons. Outgoing synapses originating from dendrites were then queried in CAVE to quantify dendrodendritic outputs.

To quantify sensory-driven triads, we analyzed EM volumes in which individual sensory subtypes were labeled. Sensory axons were manually traced as in the axoaxonic-synapse analysis. For each sensory axon, we annotated axodendritic synapses onto dendrites and classified the postsynaptic dendrite as either a vesicle-containing dendrite (VD) or a plain dendrite (PD; lacking synaptic vesicles). A triad was defined as having (i) a VD forming a dendrodendritic synapse onto a PD that (ii) also received an axodendritic synapse from the same sensory axon, with (iii) the two axodendritic contacts and the dendrodendritic contact located within a 2 μm radius. For each sensory neuron axons that we traced, triad counts were normalized to the total number of axodendritic synapses onto PDs.

### Feedforward inhibition heterotypic analysis

To characterize the postsynaptic spinal cord neurons of PSI-IhNs, PSI-IhNs were manually segmented into axonal and dendritic compartments in Neuroglancer. For axodendritic targets, synapses were queried with the axon defined as the presynaptic compartment and we identified spinal cord neurons that received at least two synapses from each PSI-IhN axon. For dendrodendritic target analysis, we screened among those that are postsynaptic to the dendritic compartments of Islet-PSI-IhN^C-HTMR/Heat^ and Islet-PSI-IhN^C-LTMR^ and receive at least 5 synapses from each PSI-IhN dendrite. Because PSI-IhN axons may also form synapses onto sensory neurons, sensory-like postsynaptic neurons were excluded. Sensory neurons were expected to have substantially more output than input synapses; therefore, neurons with a pronounced excess of output synapses were classified as sensory-like and removed. We also remove the partially reconstructed postsynaptic neurons by filtering out postsynaptic neurons with less than 400 input synapses. The spinal cord neurons receiving inputs from more than one PSI-IhN subtype were counted as postsynaptic targets of each relevant PSI-IhN population; thus, a single spinal cord neuron could belong to multiple postsynaptic neuron groups. In total, SPINE was used to analyze sensory inputs onto 1,553 postsynaptic spinal cord neurons.

For each postsynaptic neuron, we defined a homotypic selectivity index as the fraction of inputs from the corresponding sensory subtype relative to all sensory inputs. For example, for neurons postsynaptic to Islet-PSI-IhN ^C-LTMR^s, this value was calculated as the fraction of C-LTMR inputs among all sensory inputs. Postsynaptic neurons were assigned to a sensory module when a single sensory subtype accounted for more than 50% of their total sensory input, and modules were named according to the dominant sensory subtype. To calculate population-level sensory input fractions, all postsynaptic neurons associated with a given PSI-IhN population were pooled, and the fraction of total sensory input contributed by each sensory subtype was quantified.

### *Ex vivo* electrophysiological recordings

P30-90 mice were anesthetized with isoflurane and transcardially perfused with choline chloride solution consisting of (in mM): 110 Choline Cl, 2.5 KCl, 1.25 NaH_2_PO_4_, 25 NaHCO_3_, 25 glucose, 0.5 CaCl_2_, 7 MgCl_2_, 3.1 Na Pyruvate, 11.6 Na Ascorbate, 0.002 (R,S)-CPP, 0.005 NBQX, oxygenated with 95% O2 / 5% CO_2_, kept at room temperature. The lumbosacral spinal cord and attached DRGs were dissected intact and placed dorsal side up onto coverslips coated with poly-l-lysine (Sigma, P8920). Tissue was then allowed to recover for 30 minutes in ACSF consisting of (in mM): 127 NaCl, 2.5 KCl, 1.25 NaH_2_PO_4_, 1.5 CaCl_2_, 1 MgCl_2_, 26 NaHCO_3_, 25 Glucose, 0.002 (R,S)-CPP, 0.005 NBQX, kept at room temperature and oxygenated with 95% O_2_ / 5%. The coverslip was then transferred to a small chamber held at 35 °C containing 0.01–0.02% collagenase P (Sigma, 11213857001) dissolved in ACSF for 15-20 minutes before being returned to ACSF at room temperature.

The coverslip was then transferred to an upright microscope. Experiments were performed in recirculated ACSF containing (in mM): 127 NaCl, 2.5 KCl, 1.25 NaH_2_PO_4_, 1.5 CaCl_2_, 1 MgCl_2_, 26 NaHCO_3_, 25 Glucose, 0.002 (R,S)-CPP, 0.005 NBQX, kept at room temperature and oxygenated with 95% O_2_ / 5% CO_2_. A glass monopolar stimulus electrode (2-3 MΩ) was inserted into the gray matter of the dorsal horn of the spinal cord 100-400 µm rostral to the dorsal root entry of L3, L4, or L5 DRG. The corresponding DRG neurons were then visualized using DIC for random recordings (Acr1 stimulation), or subtypes were identified with fluorescent reporters (ChR2 stimulation). Borosillicate recording electrodes (9-10 MΩ) filled with internal solution consisting of (in mM): 130 K-gluconate, 3 KCl, 10 HEPES, 0.5 EGTA, 3 MgATP, 0.5 NaGTP, 5 Phosphocreatine-Tris_2_, 5 Phosphocreatine-Na_2_, pH 7.2 with KOH, were visually guided to the target DRG neuron under high positive pressure to avoid sealing onto satellite glial cells. After whole-cell recordings were established, electrical stimulation was used to establish intact connectivity between the DRG soma and its central terminal in the dorsal horn. The microscope was then positioned over the dorsal horn stimulus electrode, and optical stimuli were delivered over the dorsal horn through the microscope objective (40x objective, 473 nM, 2 ms pulse duration, 50 mW/mm^2^). For activation of Acr, the same light stimulus was used as for ChR2 activation, delivered through the objective over the dorsal horn or directly over the recording electrode. Recordings of DRG neurons were made using a Multiclamp 700B (Molecular Devices) and acquired using a Digidata 1550B with Clampex 11 software (Molecular Devices) at 20 kHz.

For identification of DRG neuron subtypes, the following mouse lines and viral tools were used: Aβ RA-LTMRs, *Np2yr-GFP*; *Vgat^IRES-Cre^; Rosa26^LSL-ChR2/+^*. For Aδ-LTMRs, *TrkB^eGFP^; Vgat^IRES-Cre^; Rosa26^LSL-ChR2/+^*; For C-LTMRs, *TH^CreER^; Vgat-ChR2* and IP injection of AAV9-FLEX-tdTomato at P0. For C-HTMRs, *Calca-GFP; Vgat-ChR2 and Mrgprd^eGFP^*; *Vgat-ChR2*. For Aδ-HTMRs, *Calca-GFP; Vgat-ChR2*.

For optical activation of Acr1 in DRG neurons, the following mouse lines and viral tools were used: *Rosa26^LSL-gtAcr1-FRED/+^*; AAVrg-Ef1a-Cre injected into hindlimb glabrous skin. C-HTMRs were identified by C-fiber conduction velocity and by the absence of T-type rebound firing following hyperpolarization. C-HTMRs also lacked the delayed firing pattern after current injection that is normally observed in C-LTMRs (Zheng et al., 2019).

### Model of heterotypic feedforward inhibition

We constructed a firing rate-based model to investigate the effect of observed heterotypic feedforward inhibitory motif on response of downstream populations in the dorsal horn. Specifically, we focused on the transformation of force tuning curves in going from sensory neurons to corresponding excitatory neurons downstream. The model consists of three sensory subtypes: light touch (C-LTMR), medium-force touch (MrgD^+^ C-HTMR/Heat), and high-force touch (A*δ*-HTMR) sensory neurons, each with a corresponding excitatory and inhibitory PSI-IhN units (**Figure S13c,d,e**). In the model, each population is represented by a single firing-rate unit. Sensory afferent firing rates are modelled as ℎ*_c_*(*F*) = [*s_c_* tanh(*g_c_*(*F* − *θ_c_*)/*s_c_*]_+_, where *F* represents mechanical force (in milli-Newtons), [⋅]_+_ represents rectification at zero, *s_c_*, *g_c_*, and *θ_c_* denote the saturation rate, gain, and force threshold of the *c^t^*^ℎ^ sensory subtype, respectively. These three parameters were set based on functional experiments of (Lezgiyeva et al., 2026; Ma et al., 2012; Pitcher et al., 2016; Qi et al., 2024): the threshold parameters were set to *θ_L_* = 5, *θ_M_* = 20, *θ_H_* = 50 *mN*, with a common gain of *g* = 3, and saturation rates of *s_L_* = 35, *s_M_* = 30, *s_H_* = 25 *Hz* (*L*, *M*, and *H* subscripts denote light-, medium-, and high-force sensory subtypes, respectively). The dynamics of the model takes the form *τ dr_a_*⁄*dt* = −*r_a_* + [∑*_b_ J_ab_ r_b_* + ∑*_c_ W_ac_* ℎ*_c_*]_+_, where *r_a_* is the firing rate of the *a^t^*^ℎ^ dorsal horn units, namely, C-LTMR module, C-HTMR/Heat module or A*δ*-HTMR module, which form the excitatory units, and their corresponding PSI-IhN subtypes. The time constant was set to *τ* = 1. *W* contains the feedforward weights from sensory afferents (S) to dorsal horn units: these weights onto their matched excitatory (E) and PSI-IhN (I) interneuron units were set to *β_ES_* = 1, and *β_IS_* = 3, respectively. Lateral inhibition was implemented such that each PSI-IhN suppressed both its matched excitatory unit (homotypic FFI, *β*^homo^ = 0.1) and the adjacent medium-force excitatory unit (heterotypic FFI, *β*^hetero^ = 0.1). The medium-force PSI-IhN unit, however, only inhibited its matched excitatory unit (**Figure S13c**).

## Acknowledgements

The authors thank T. Docter, A. Shuster, S. Deng, S. Markman, K. Ng and the members of the Ginty and Lee laboratories for discussions and comments on the manuscript; J. Joroff, E. Yacharn, E. Xue, K. Ohgami, S. Simhadri for tracing neurons; S. Gerhard for advice on analyzing neuron skeletons; F. Collman and C.S.-Mizell for advice analyzing neuron mesh data; J. Phelps, A.T. Kuan, D.G.C. Hildebrand, B. Graham, and K. Kris for advice on EM imaging and alignment; R. Schalek for performing micro-CT scans on EM samples; HMS IT staff for assisting with data storage; L. Qi for assistance with animal surgery and breeding. The authors gratefully acknowledge BossDB.org for providing data storage and access services that support the sharing of the datasets associated with this publication (R24MH114785, Hider et. al 2022 – https://doi.org/10.3389/fninf.2022.828787). This work was supported by Stuart H.Q. and Victoria Quan fellowships (W.X., Q.Z. and J.L.), NIH grant R35 5R35NS132196 (D.D.G.), the K. Lisa Yang Brain Body Center at Harvard Medical School (D.D.G.), the Edward R. and Anne G. Lefler Center for Neurodegenerative Disorders (D.D.G.) and the R&D Impact Foundation Neurobiology Spark Grant (W.C.A.L.). D.D.G. is an investigator of the Howard Hughes Medical Institute.

## Author contributions

W.X., K.D., Q.Z., D.L.P., W.-C.A.L. and D.D.G. conceptualized the project and designed experiments. W.X. generated the EM datasets with help from M.L. to set up the microscope. K.D. and W.X. developed and applied the SPINE model with input from Q.Z. and S.D. J.T. designed and performed the *ex vivo* physiology experiments. T.N., T.M. and N.K. developed and applied the segmentation pipeline. J.A.B. developed and applied the synapse prediction pipeline. A.H. set up CAVE for reconstruction. W.X., S.C., Q.Z., B.M.L., M.F.M., J.N., R.B., A.S., L.S.C., A.N. and J.L. reconstructed the neurons. W.X., K.D., A.H., analyzed the connectivity data. R.K. performed modeling on feedforward inhibition. W.X., W.-C.A.L., and D.D.G. wrote the manuscript with input from other authors.

## Data availability

The volumetric EM datasets generated in this study, including the dSC1 dataset and dSC_APEX datasets, segmentation, and cell-type annotations are open source and publicly available through: https://github.com/htem/spinal_cord_connectome.

## Code availability

Code for the development and application of SPINE is available at: https://github.com/htem/dorsalhorn-ml. Code for analysis and figure generation is available at: https://github.com/htem/spinal_cord_connectome.

